# Barley guanine nucleotide exchange factor *Hv*GEF14 is an activator of the susceptibility factor *Hv*RACB and supports host cell entry by *Blumeria graminis* f.sp. *hordei*

**DOI:** 10.1101/2021.08.04.455079

**Authors:** Adriana Trutzenberg, Stefan Engelhardt, Ralph Hückelhoven

## Abstract

In barley (*Hordeum vulgare*), the function of ROPs appears central to polar cell development and the interaction outcome with parasitic fungi but little is known about ROP activation. Guanine nucleotide exchange factors (GEFs) facilitate the exchange of GDP with GTP and thereby turn ROPs into a signalling-activated ROP-GTP state. Plants possess a unique class of GEFs harbouring a plant specific ROP nucleotide exchanger domain (PRONE). We performed phylogenetic analyses and annotated barley PRONE GEFs. *HvGEF14* is expressed in leaf epidermal tissue and downregulated after inoculation with *Blumeria graminis* f.sp. *hordei*. Protein-protein interaction assays indicate that *Hv*GEF14 and the type I barley ROP protein *Hv*RACB can interact in yeast and *in planta*. Overexpression of *Hv*GEF14 further recruited the ROP-GTP downstream interactor *Hv*RIC171 to the cell periphery and let to interaction of *Hv*RACB with an extended CRIB (Cdc42/Rac Interactive Binding motif) peptide of *Hv*RIC171 in a similar manner as constitutively activated *Hv*RACB. Finally, the over expression of *HvGEF14* led to enhanced susceptibility to fungal entry while *HvGEF14* RNAi provoked a trend to more penetration resistance. *Hv*GEF14 might therefore play a role in the activation of *Hv*RACB in barley epidermal cells, which is crucial for fungal penetration success.

**Highlight:** The activated GTPase *Hv*RACB is a susceptibility factor of barley in response to *Blumeria graminis* f.sp. *hordei*. The newly discovered guanine nucleotide exchange factor *Hv*GEF14 is a *Hv*RACB activator.

## Introduction

In plants, cell polarity is central to cellular development and responses to external stimuli. ROPs (RHO of plants) are small monomeric GTPases that function as signalling hubs regulating a multitude of cell polarity processes that often involve cytoskeleton reorganization (Mucha *et al*., 2011). Pollen tube growth, the development of epidermal pavement cells and root hairs, but also processes that are vital during plant-microbe interactions are examples for ROP-regulated processes (Engelhardt *et al*., 2020; Zheng and Yang, 2000). ROPs are considered molecular switches due to their ability to shuttle between a signalling-inactive, GDP-bound state, and a signalling-activated GTP-bound state (Bloch and Yalovsky, 2013). An interaction with downstream signalling partners, and therefore signal transduction, only occurs in the GTP-bound state (Nagawa, Xu, and Yang 2010). Certain point mutations can lock ROPs in either the GTP-bound (constitutively activated (CA)) or GDP-bound (dominant negative (DN)) conformation. Manipulating ROP downstream signalling in such a way can lead to strong effects in a variety of polar growth processes. For example, *Arabidopsis thaliana* ROP6^DN^ expressing plants show severe developmental phenotypes as well as misshaping of pavement cells, indicating the importance of ROP-regulated cell polarity (Poraty-Gavra *et al*., 2013).

To locally interact with downstream signalling partners, ROPs require to be membrane-associated, which is achieved by posttranslational lipid modifications. Based on the primary sequence in the C-terminal part, ROPs can be divided into two groups that both undergo distinct lipidations: type I ROPs contain a CaaL-prenylation motif, whereas type II ROPs are S-acylated via a GC-CG box. Type I ROPs can also be additionally S-acylated in an activation dependent manner (Winge *et al*., 2000; Yalovsky, 2015). Amongst others, ROP signalling targets the cytoskeleton (Chen and Friml, 2014). In barley, the activation of ROP-signalling influences the organization of actin filaments as well as the organization of microtubules via the interaction of activated ROPs with scaffold proteins and **M**icrotubule-**A**ssociated ROP **GAP 1**, *Hv*MAGAP1 (Hoefle *et al*., 2020; Opalski *et al*., 2005).

Barley (*Hordeum vulgare* L. (*Hv*)) contains six ROPs, which include *Hv*RACB and *Hv*RACD of type I and *Hv*RAC1, *Hv*RAC3, *Hv*ROP4, *Hv*ROP6 of type II (Schultheiss *et al*., 2003). Studies using stable transgenic barley plants overexpressing constitutively activated variants of either *Hv*RACB, *Hv*RAC1 or *Hv*RAC3 indicated a role for these ROPs in root and shoot development (Pathuri *et al*., 2008; Schultheiss *et al*., 2005). Transgenic plants are also characterized by increased epidermal cell expansion and failure of polar growth in root hairs. Additionally, different susceptibility levels during the interaction with fungal leaf pathogens such as *Blumeria graminis* f.sp. *hordei* (*Bgh*) and *Magnaporthe oryzae* suggest an involvement during fungal penetration (Pathuri *et al*., 2008).

*Hv*RACB in particular has been studied in some detail for its function in susceptibility to penetration and accommodation of haustoria from *Bgh* (Engelhardt *et al*., 2020). Stable transgenic *Hv*RACB RNAi plants with reduced expression levels of *HvRACB* do not develop root hairs and produce less and misshaped stomatal subsidiary cells, which normally result from polar cell division. At the same time, both stable and transient silencing of *HvRACB* limits fungal penetration success. Hence, *Hv*RACB is considered a key developmental protein that is co-opted by *Bgh* during pathogenesis (Scheler *et al*., 2016; Schultheiss *et al*., 2002). *Hv*RAC1 has also been implicated in the generation of membrane domains enriched with the ROP-scaffolding protein *Hv*RIPa at sites of fungal attack. The two proteins interact at the plasma membrane, and together with *Hv*MAGAP1 they provoke membrane asymmetry and microtubule reorganization in the cell cortex (Hoefle *et al*., 2020).

Due to the vast number of signalling processes ROPs are involved in, it is apparent that tight regulation of these molecular switches is required to fine-tune the complex cellular mechanisms following ROP activation. This regulation is largely controlled by three classes of regulatory proteins: **G**uanine nucleotide **E**xchange **F**actors (GEF), **G**TPase **A**ctivating **P**roteins (GAP) and **G**uanine nucleotide **D**issociation **I**nhibitors (GDI) (Nagawa *et al*., 2010; Vetter and Wittinghofer, 2001; Zheng and Yang, 2000). Besides downstream signalling interactors, activated ROPs are also targeted by GAPs, which abolish ROP-GTP signalling by enhancing the intrinsically low GTPase activity. This leads to GTP hydrolysis and therefore to switching to the GDP-bound, signalling-inactive state (Wu *et al*., 2000). An example of this is *Hv*MAGAP1 which interacts with *Hv*RACB and *Hv*RAC1. The presumable negative regulation of *Hv*RACB signalling activity by over-expression of *Hv*MAGAP1 correlates with limiting *Bgh* penetration success (Hoefle *et al*., 2011).

GDIs prevent GDP-bound ROPs from membrane association and sequester them in the cytosol by masking the C-terminal prenyl modification, thereby precluding an interaction with downstream interaction partners (Nibau *et al*., 2006). Regarding ROP signalling, GEFs are considered ROP-signalling activating proteins that interact with GDP-bound ROPs to induce a conformational change facilitating the exchange of GDP with highly abundant GTP (Berken *et al*., 2005; Thomas *et al*., 2007; Thomas *et al*., 2009).

ROP-signalling activating GEFs can be divided into sub-groups based on their protein domain structures and primary sequence. Mammalian GEFs possess two types of conserved functional domains: **d**iffuse B-cell lymphoma **h**omology – **p**leckstrin **h**omology (DH-PH) and **D**OCK **h**omology **r**egion **2** (DHR2) (Schmidt and Hall, 2002). In plants, there has only been one example of DH-PH GEFs and several reports on DHR2-type GEFs. The *O. sativa* DH-PH GEFs *Os*SWAP70A and B interact with *Os*RAC1 and perform nucleotide exchange via their DH domain in order to facilitate defence-related signalling in response to chitin elicitation (Yamaguchi *et al*., 2012). The same group also confirmed the corresponding *A. thaliana* homolog *At*SWAP70 to be involved in plant defence signalling during effector and pattern triggered immunity (Yamaguchi and Kawasaki, 2012).

An example of a DHR2 GEF in plants is the so called SPIKE-GEF which has initially been identified by a forward genetic screen of *A. thaliana* mutants, in which the *spike1* mutant showed severe developmental defects, including misshaped trichomes (Qiu *et al*., 2002). More recently, *Lj*SPIKE1 has been found to activate *Lj*ROP6 during the polar growth of infection threads in root hairs of the legume *Lotus japonicus* (*Lj*) (Liu *et al*., 2020).

In addition, plants have evolved a specific class of GEFs with a highly conserved **P**lant specific Rac/**RO**p **N**ucleotide **E**xchanger (PRONE) domain (Berken *et al*., 2005; Gu *et al*., 2006). The family of PRONE-GEFs in the model plant *A. thaliana* consists of 14 members that have been studied for their function in various polar growth processes, plant development (Chang *et al*., 2013; Chen *et al*., 2011; Gu *et al*., 2006; Huang *et al*., 2018a) and immunity (Qu *et al*., 2017). So far, the PRONE domain is the only identified conserved part of these proteins. This domain can be divided into three subdomains that show a high level of sequence conservation. The PRONE domain contains several interfaces that were shown to physically interact with ROPs in a heterotetrameric complex of two ROPs with two PRONE-GEFs. After binding to ROPs, the PRONE domain facilitates a rearrangement of the GTPase which leads to GDP release. Nucleotide-free ROPs and GEFs can further interact as a stable complex (Berken *et al*., 2005; Chang *et al*., 2013; Gu *et al*., 2006; Thomas *et al*., 2007). The PRONE domain is flanked by N- and C-terminal variable regions that likely possess regulatory functions. Especially the C-terminal regulatory stretch is subject to phosphorylation by upstream **R**eceptor **L**ike **K**inases (RLK) that have been implicated in developmental as well as immunity related signalling pathways (Fehér and Lajkó, 2015). For instance, the C-terminus of *At*GEF12 has been reported to inhibit GEF function and *At*GEF12 interacts with ROPs only after phosphorylation by the pollen RLK *At*PRK2 (Zhang and McCormick, 2007). Another example of this interaction is the multifunctional RLK FERONIA (FER) that interacts directly with *At*GEF14 in *A. thaliana*. Further downstream, *At*GEF14 interacts with *At*ROP6 which functions in polar growth of epidermal cells (Lin *et al*., 2018) and together with *At*RIC1 facilitates microtubule organization in order to deal with mechanical stress (Tang *et al*., 2021). In barley, a number of RLKs are involved in susceptibility towards *Bgh* and their transcript levels are partially influenced by the abundance and the activation status of *Hv*RACB (Schnepf *et al*., 2018).

Furthermore, *At*GEF14 was shown to localize to the apical region of pollen tubes and interacts with *At*ROP1, a regulator of pollen tube growth (Gu *et al*., 2006). In roots, *At*GEF14 may be also involved in polar growth processes. It accumulates at the root hair initiation site before being replaced by other GEFs during the initiation and elongation phase (Denninger *et al*., 2019). Based on these first reports of PRONE GEFs in *A. thaliana*, plant specific GEFs have been identified in many other species, including economically important crops like *Oryza sativa*, in which six of the 11 putative PRONE-GEFs have been studied in some detail. As in *A. thaliana*, *O. sativa* GEFs are also important in floral organ development. *Os*GEF7B (*Os*GEF2 in other publications, see (Akamatsu *et al*., 2015; Akamatsu *et al*., 2013)), for example, has been shown to be an important regulator of the seed setting rate by regulating downstream *Os*RAC GTPases (Huang *et al*., 2018b). Moreover, four pollen-specifically expressed GEFs have been reported with distinct localisations during the development of pollen tubes (Kim *et al*., 2020). *Os*GEF10 has been found to activate *Os*RAC1 in order to form small cuticular papillae and to regulate early plant development (Yoo *et al*., 2011). In addition, *Os*GEF1 interacts with OsCERK1, a central part of the rice chitin receptor complex, and activates *Os*RAC1 for its function in defence of *M. oryzae* (Akamatsu *et al*., 2013).

This work investigates barley PRONE-GEFs, focussing on *Hv*GEF14. We report that *Hv*GEF14 is expressed in the leaf epidermis and downregulated after inoculation with *Bgh*. *Hv*GEF14 can bind barley ROPs in yeast and *in planta* and may be involved in activation of disease susceptibility-associated barley ROPs in leaf epidermal cells.

## Materials and Methods

### Plant and pathogen propagation and maintenance

Spring barley (*Hordeum vulgare, Hv*) cultivar Golden Promise (propagated in greenhouse) was grown in a climate chamber at 20 °C, 50 % humidity and long-day conditions (16h light, 8h dark) for 7-8 days in standard potting soil (Einheitserde Classic). *Blumeria graminis* f.sp. *hordei* race A6 (*Bgh*) was maintained on 7-21 days old *H. vulgare* cv. Golden Promise (propagated in field conditions) in a climate chamber with 18 °C and 65 % humidity in long day conditions (16h light, 8h dark).

### Cloning procedure

Open reading frames of *Hordeum vulgare* cv. Golden Promise genes were amplified from whole leaf cDNA with primers (Tab. 5) designed based on sequences from the Barley Genome (International Barley Genome Sequencing Consortium (2012)). Constructs were cloned into destination vectors (Tab. 4) using BP and LR clonase following manufacturer’s instructions (Invitrogen). Plasmids were prepared via column purification (Machery Nagel).

### ML analysis and phylogenetic tree construction

Muscle alignment of 14 *Arabidopsis thaliana*, 11 *Oryza sativa* (sequences downloaded from TAIR and NCBI on 10.05.2021) and 11 *H. vulgare* PRONE-GEFs (MOREX genome version 3) was performed in SeaView Software. A maximum likelihood (PhyML) analysis was performed with an LG model, bootstraps with 100 replicate, model-given amino acid equilibrium frequencies, NNI tree searching and five random starts. *O. sativa* PRONE-GEFs were annotated based on the *A. thaliana* nomenclature and *H. vulgare* PRONE-GEFs were annotated according to their homologues in *O. sativa*. The resulting TBE tree’s design was further adjusted in InkScape.

### RNA extraction and cDNA synthesis

Plant material from seven days old *H. vulgare* cv. Golden Promise was collected. In three biological replicates, whole leaf and epidermal peels were cut and directly frozen in liquid N_2_. Leaf tissue was directly ground in a tissue lyser with glass beads. RNA was extracted with TRIzol according to the protocol in (Chomczynski and Sachhi, 1987) and DNAseI digestion was performed. Total RNA was visualized on 1% agarose gel and the concentration measured via spectrophotometer (NanoDrop). Subsequently, cDNA synthesis was performed from 1 μg RNA with the QuantiTect Reverse Transcription Kit (Quiagen) according to the supplier’s protocol. cDNA from 1 μg RNA was diluted 1/10 for further analysis and stored at -20 °C. cDNA quality was controlled on 1% agarose gel and visualised with Ethidium-Bromide staining under UV-excitation.

### qRT-PCR

Appropriate primers for *HvGEF14* (HORVU6Hr1G068660.3, Tab. 5) were designed with Primer3Plus (www.bioinformatics.nl/cgi-bin/primer3plus/primer3plus.cgi), secondary structure of primers was checked with Thermo Multiple Primer Analyser (https://www.thermofisher.com/de/de/home/brands/thermo-scientific/molecular-biology/molecular-biology-learning-center/molecular-biology-resource-library/thermo-scientific-web-tools/multiple-primer-analyzer.html) and specificity of the amplicon was tested via BLAST. Primer efficiency and amplification factor was analysed by a standard curve of a three-step dilution series. A calibration curve was calculated with -3.116 slope, r^2^ of 0.951 and y-intercept at 29.58. The power was determined via the online calculator “qPCR Efficiency Calculator” (Thermo Fischer). 10 μl reactions were prepared with the Maxima 2xSYBR Green/ROX qPCR Master Mix (Thermo Scientific), primers were added at 200nM concentration and 1 μl of 1/10 cDNA dilution was used. qRT-PCR was run on Aria Mx3000 thermocycler (Agilent) with 40 cycles at 60°C annealing temperature for 10 seconds and 72°C elongation for 15 seconds and subsequent melting curve (65°C – 95°C) for quality control. Measurements with melting curve peaks varying from all other samples were excluded and a maximum difference of 0.5 Cq between technical replicates was allowed. There were no Cq values detectable for NTC of the housekeeping gene and for *HvGEF14* a Cq of 36.67 was measured for NTC. *HvUbiquitin* was measured as a housekeeping gene in all runs due to its reliable Cq values in all samples that did not differ more than 2 Cq and successful usage in previous work (Schnepf *et al*., 2018). Threshold values were exported from the AriaMx software (Agilent) and foldchanges were calculated via the 2^−ΔΔCt^ method by Livak and Schmittgen (2001).

### Yeast-2-Hybrid

Protein-protein interactions via yeast-two-hybrid (Y2H) assays were performed as described in the Matchmaker protocol (Clontech). *Hv*RACB (amino acids 1-193) and *Hv*RAC1 (amino acids 1-211) open reading frames were cloned with a premature STOP codon to express truncated ROP proteins lacking their C-terminal prenylation/acylation signal sequence to avoid membrane localisation in yeast cells. *Saccharomyces cerevisiae* strain AH109 was transformed with pGBKT7 and pGADT7 plasmids (Tab. 4) containing the specific gene of interest and cultivated for 3-6 days at 30°C on synthetic dropout medium lacking amino acids leucine and tryptophan (SD-L-W) medium. 5ml of liquid SD-L-W medium was inoculated with yeast colonies and, upon overnight cultivation at 30°C, dilutions were dropped on SD-L-W and SD-L-W-H (SD lacking amino acids leucine, tryptophan, histidine) or SD-L-W-H-Ade (SD lacking amino acids leucine, tryptophan, histidine and nucleotide adenosine) plates and incubated at 30°C until distinct colony growth.

### Protein extraction from yeast and *N. benthamiana*

Transformed yeast was cultured in 4 ml synthetic dropout medium lacking amino acids leucine and tryptophane (SD-L-W) overnight and centrifuged at 4000g for 5 min at 4°C. Cells were washed in 100 μl 2 M LiAc and subsequently incubated for 5 min at room temperature in 100 μl 0.4 M NaOH. Pellets were collected and 50 μl 4x SDS-sample buffer was added. After vortexing, samples were boiled at 95 °C for 5 min and shortly centrifuged before loading onto an SDS-gel (Zhang *et al*., 2011).

3 leaf discs (12 mm tissue punch) were collected from *A. tumefaciens*-transformed *N. benthamiana* leaves 48 hpt and directly frozen in liquid N_2_. The frozen plant material was homogenised in a tissue lyser with glass beads and 200 μl 4x SDS sample buffer was added. After vortexing, samples were boiled at 95°C for 10 min and shortly centrifuged before loading onto an SDS-gel.

### SDS-PAGE and Western Blot

Extracted proteins were separated by electrophoresis in a 12% acrylamide gel and blotted onto PVDF membrane via semi-dry Western Blotting (protein extracted from yeast) or wet Western Blotting (protein extracted from *N. benthamiana*). The membrane was blocked with 5% milk in PBS buffer and incubated with specific antibodies (see figure legends). Proteins were detected by chemiluminescence with SuperSignalTM West Dura or FEMTO Chemiluminescence-Substrate (Thermo ScientificTM).

### Transient biolistic transformation of barley epidermal cells

Barley cv. Golden Promise seven days old detached leaves were transformed by particle bombardment as described previously (McCollum *et al*., 2020). 2 μg of plasmid/transformation was used in FRET-FLIM experiments. For fungal penetration efficiency experiments, 1 μg plasmid/transformation for the gene of interest and 0.5 μg plasmid/transformation of transformation marker (pUbi_GUSplus, Addgene, encoding ß-Glucuronidase) was applied.

### A.tumefaciens infiltration of N. benthamiana leaves

*Agrobacterium tumefaciens* strain GV3101 carrying binary expression vectors pGWB containing either meGFP-*Hv*RACB WT, meGFP-*Hv*RACB G15V (CA), GST-mCherry, *Hv*CRIB46-mCherry or 3xHA-*Hv*GEF14 were infiltrated into five-week-old *N. benthamiana* leaves according to Yang *et al*. (2000). Bacterial liquid cultures were grown to OD600 0.5 and mixed in equal amounts including P19 silencing suppressor. 48 hours after infiltration, proteins were extracted for Western Blotting and FRET-FLIM measurements were performed on leaf discs.

### FRET-FLIM

For protein-protein interaction studies*, Hv*GEF14 was N-terminally tagged with monomeric enhanced GFP as donor and N-terminal fusions with mCherry of *Hv*RACB variants were used as acceptors. To test *Hv*RACB activation status in planta, binary vectors of meGFP-*Hv*RACB WT or meGFP-*Hv*RACB CA were co-expressed with GST-mCherry or *Hv*CRIB46-mCherry with or without co-expression of 3HA-*Hv*GEF14 via *A. tumefaciens* infiltration in *N. benthamiana* leaves.

Microscopy of transiently transformed *H. vulgare* epidermis cells and *A. tumefaciens*-infiltrated *N. benthamiana* leave discs was performed with an Olympus FV 3000 microscope with 488 nm (20 mW) and 561 nm (50 mW) diode lasers. GFP photons were excited with a 485 nm (LDH-D-C-485) pulsed diode laser and time-correlated single photon counting (TCSPC) was performed with 2x PMA Hybrid 40 photon counting detectors. A minimum of 1000 photon counts were collected and subsequently analysed with the PicoQuant SymPhoTime 64 software. N-exponential reconvolution and decay curve fitting with daily measured or calculated IRF, for *H. vulgare* and *N. benthamiana* respectively, was applied to gain a fit with X^2^ values between 0.9 and 1.2.

### Confocal microscopy

Transiently transformed barley epidermis cells were imaged 24 hours post transformation (hpt) with a Leica TCS SP5 confocal microscope with hybrid HyD and PMT detectors. mCherry fluorophores were excited with a 561nm laser and detected at 570-610nm. GFP fluorescence was excited with a 488nm Argon laser and detected between 500-550nm.

### RNAi efficiency

GFP tagged *Hv*GEF14 was transiently over expressed together with overexpression of cytosolic mCherry as a transformation marker. In addition, either the empty vector or a *HvGEF14* RNAi hairpin construct of amino acids 201-401 was co-transformed and fluorescence intensity of GFP and mCherry was measured in the z-stack of confocal images taken 48 hpt. RNAi efficiency was determined by the ratio of mean mCherry-normalised GFP fluorescence in *HvGEF14* silenced cells divided by mCherry-normalised GFP-*Hv*GEF14 expressing control cells.

### Fungal penetration efficiency assays

Detached barley leaves were fixed on 0.8% H_2_O-agar and inoculated 24 hours after transient transformation with over-expression constructs or 48 hours after transient transformation with RNAi constructs with roughly 100 *Bgh* conidiospores per mm^2^. Inoculated leaves were incubated for 48 hours in a climate chamber with 18-22 °C under long day conditions (16h light, 8h dark). Inoculated leaves were stained in X-Gluc (5-Bromo-4-chloro-3-indolyl ß-D-glucuronic acid) solution 48 hours after inoculation and fixed in 80% EtOH. Fungal penetration efficiency (PE) was determined with light microscopy as described in Hückelhoven *et al*. (2003).

### Statistical analysis

Statistical analyses were performed in Rstudio. Global comparison for non-parametric data was assessed with Kruskal.test() and pairwise comparisons of non-parametric data were calculated via wilcox.test() with bonferroni p-value adjustment. Outlier tests were performed with grubbs.test(). Figures were prepared with RStudio’s ggplot2 package and adjusted in InkScape.

## Results

### Phylogenetic analysis reveals three distinct clades of barley PRONE-GEFs

To investigate guanine nucleotide exchange factors in barley, we concentrated on the well described PRONE domain-containing class of exchange factors first introduced by Berken *et al*. (2005). In total, we identified 11 PRONE domain-encoding genes in *Hordeum vulgare* cv. Morex genome version 3 (Mascher *et al*., 2021). Using the MUSCLE algorithm, we aligned full-length primary amino acid sequences of these 11 barley PRONE-GEFs (Tab. 2) with 11 PRONE-GEFs of *Oryza sativa*, and 14 PRONE-GEFs of *Arabidopsis thaliana*. Since the *O. sativa* PRONE-GEFs have not all been annotated so far, we first constructed a maximum-likelihood phylogenetic tree via PhyML, in which all *O. sativa* PRONE-GEFs were annotated according to their primary sequence similarity to *A. thaliana* PRONE-GEFs using the nomenclature by Berken *et al*. (2005) (Fig. S1). Subsequently, another PhyML calculation was performed to obtain the phylogenetic tree with all three species (Fig. 1). *H. vulgare* PRONE-GEFs were then annotated according to the most closely related *O. sativa* PRONE-GEFs to reach a nomenclature that is consistent for grasses. That way, the barley PRONE-GEFs were named *Hv*GEFs 1, 2, 3a, 3b, 3c, 7, 9a, 9b, 9c, 10 and 14.

**Fig. 1:**
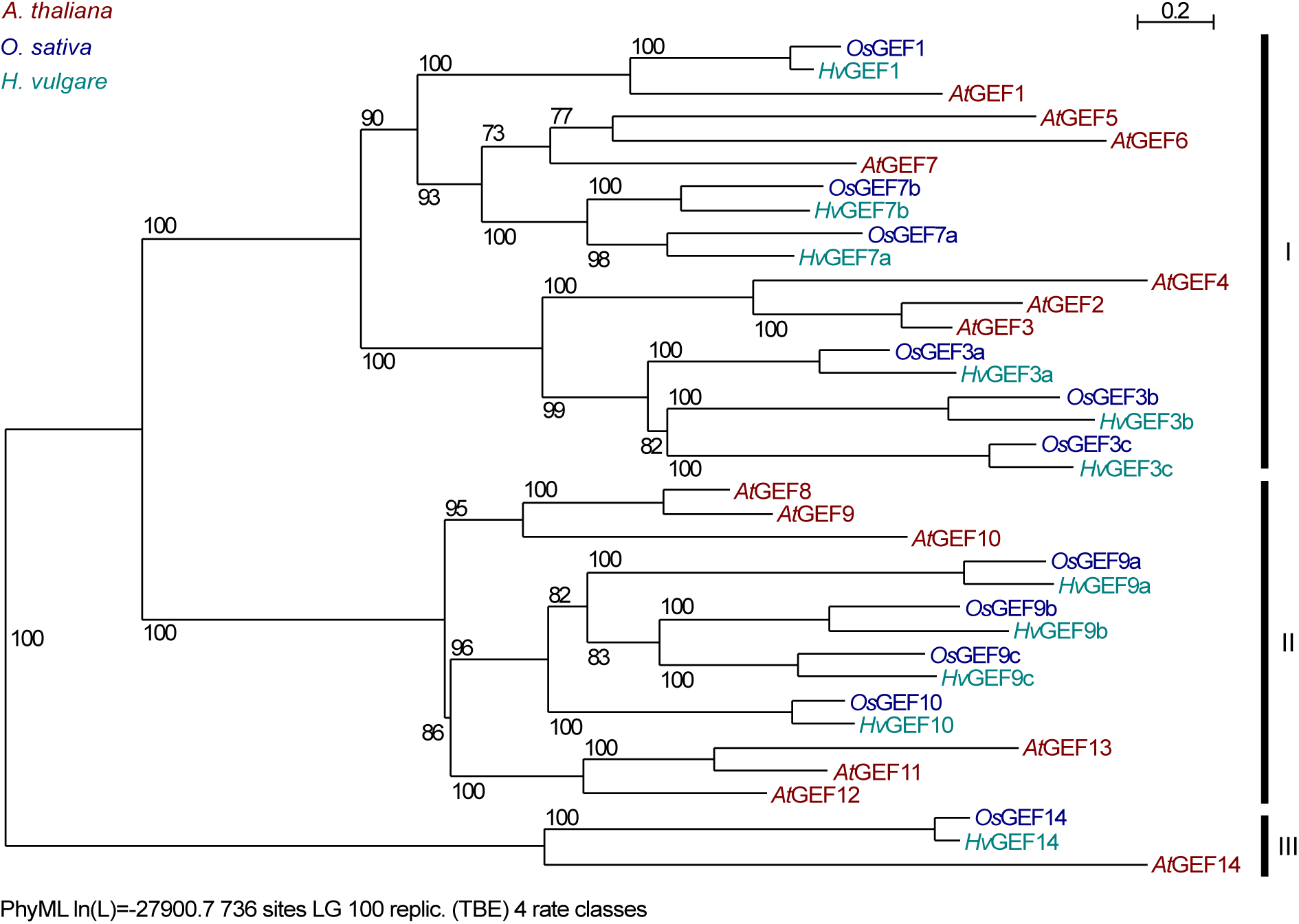
PRONE-GEFs cluster in three distinctive clades. Phylogenetic analysis of PRONE-GEFs of *Arabidopsis thaliana* (*At*), *Oryza sativa* (*Os*) and *Hordeum vulgare* (*Hv*) based on MUSCLE alignment. Maximum likelihood tree was calculated based on available amino acid sequences (NCBI and barley Morex genome version3) and annotation of barley PRONE-GEFs was performed based on this tree.

The resulting phylogenetic tree shows three distinct groups: clade (I), including PRONE-GEFs 1-7, clade (II) clustering PRONE-GEFs 8-13 and a third clade consisting of only PRONE-GEF14 proteins from the three different species (Fig. 1). Notably, in clades (I) and (II) we specifically found a clustering of PRONE-GEFs from both monocot species and many high confidence branches with only *A. thaliana* PRONE-GEFs suggesting PRONE-GEF gene duplications after the separation of monocots from dicots. The three PRONE-GEF14 sequences differ not only in their variable N- and C-terminal regions from the other PRONE-GEFs but present higher levels of amino acid changes in the PRONE domain itself when compared to all other PRONE-GEFs (Fig. S1, Tab. S1). PRONE-GEF14 proteins are not present in the moss *Physcomitrium patens* or the lycophyte *Selaginella moellendorffii* (Eklund *et al*., 2010). Despite its unique position in the phylogenetic tree, the overall structure of the *Hv*GEF14 PRONE domain with its three sub-domains and essential amino acids for GEF-GEF and GEF-ROP interactions is conserved when compared to the other barley PRONE-GEFs (Fig. 2).

**Fig. 2:**
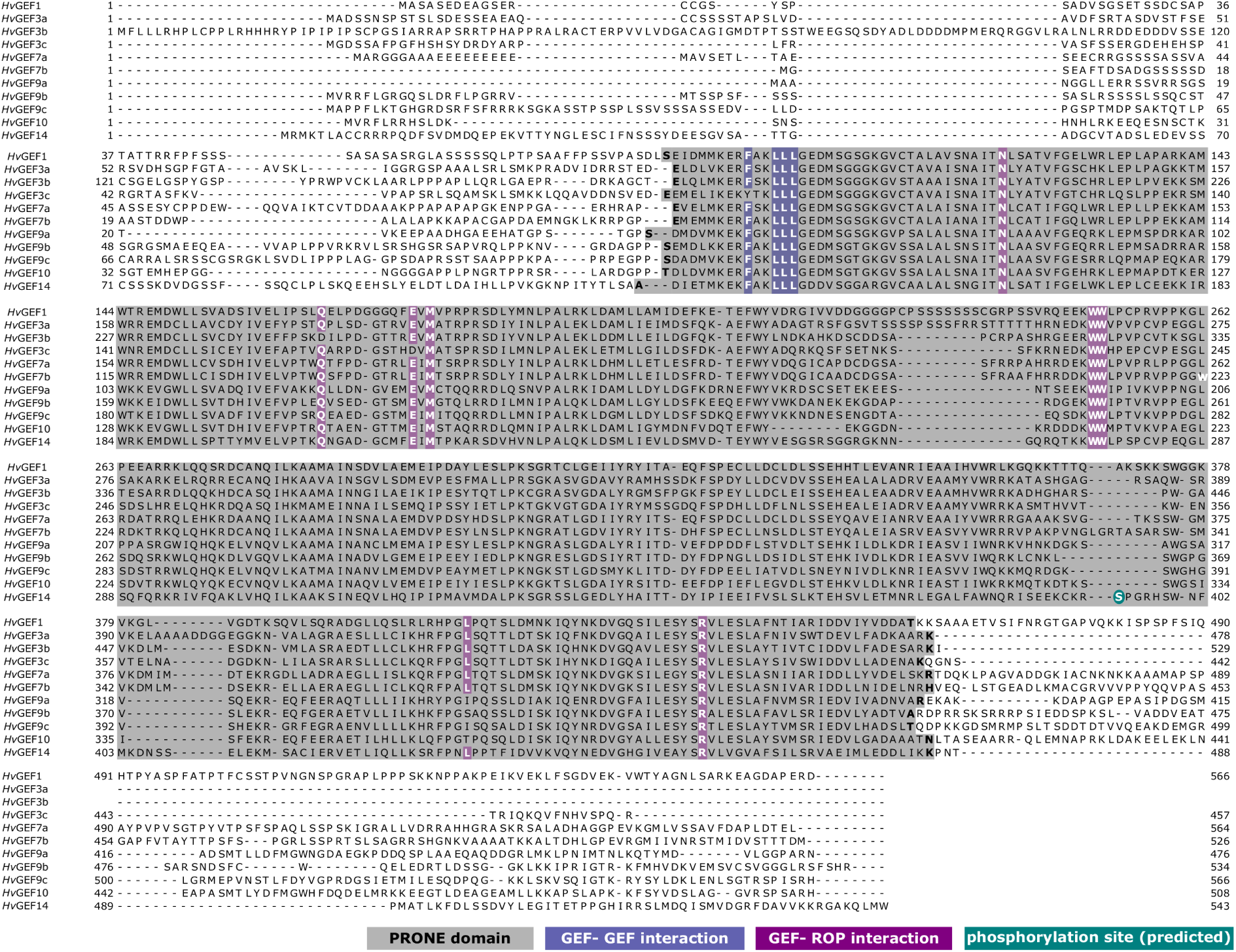
Hordeum *vulgare* (*Hv*) PRONE-GEF. MUSCLE alignment with annotation of PRONE domain (according to NCBI prediction) in grey, GEF-GEF interaction residues and ROP-GEF interaction residues highlighted in purple. Predicted phosphorylation site in *Hv*GEF14 highlighted in turquoise. Protein IDs according to NCBI accessions and annotation based on MUSCLE alignment with *O. sativa* PRONE-GEFs (see Fig. S1)

The MUSCLE alignment of 11 barley PRONE-GEFs highlights the predicted PRONE domains (see Fig. 2) with varying length of 344 amino acids (*Hv*GEF10) to 379 amino acids (*Hv*GEF3a) (Fig. S1, Tab. S2). *Hv*GEF3a has no variable C-terminal region beyond the PRONE domain, and the C-terminus of *Hv*GEF3b and *Hv*GEF3c are comparably short (Fig. 2, Fig. S1, Tab. S2). *Hv*GEF9a is predicted to contain only a short variable N-terminus of 43 amino acids. It is evident from the alignment that, regarding the primary sequence, *Hv*GEF14 substantially differs most from all other barley PRONE-GEFs. In addition, the variability in primary sequence of *Hv*GEF14 is highest when compared to the consensus sequence of all 11 *Hv*PRONE-GEFs (Fig. S1, Tab. S1). However, important residues for GEF-GEF interaction (Thomas *et al*., 2007) such as phenylalanine 133 and leucine 138 (in *Hv*GEF14) as well as residues involved in GEF-ROP interaction N161, Q206, E215, M217, W275, W276, L434, and R460 (Thomas *et al*., 2007) are conserved in *Hv*GEF14 (Fig. 2). Furthermore, *Hv*GEF14 full length and PRONE domains only were shown to dimerise in Y2H (Fig. S3). Interestingly, serine 394 in the *Hv*GEF14 PRONE domain, is a predicted phosphorylation site, based on mass spectrometry analysis of phosphorylation sites in *A. thaliana* GEF14 (Mergner *et al*., 2020). This putative phosphorylation site is conserved in *At*GEF14 and *Hv*GEF14 but not outside the GEF14 clade.

### *Hv*GEF14 is expressed in epidermal cells and downregulated after *Bgh* inoculation

Based on their gene expression, *Arabidopsis* PRONE-GEFs have been shown to operate in different plant tissues (Shin *et al*., 2009; Zhang and McCormick, 2007). To understand the potential function of barley *PRONE-GEF*s, gene expression patterns in different tissues were investigated. An initial *in silico* expression analysis of the eight barley *PRONE-GEFs* was performed by consulting the RNAseq database provided by The James-Hutton-Institute (https://ics.hutton.ac.uk/barleyGenes). In accordance with previous studies in *A. thaliana*, the RNAseq data shows that some of the barley *PRONE-GEF*s were expressed in distinct tissues only. *HvGEF3a*, for example, is mainly expressed in the developing embryo. Other barley *GEFs*, like *HvGEF1*, show a broader expression pattern with highest number of RNA fragments detected in embryos, root and grains. *HvGEF14* is the most ubiquitously expressed PRONE-GEF of barley with highest fragment counts in almost all tissues, except in senescing leaves. Interestingly, *HvGEF1* and *HvGEF14* are the only GEF genes expressed in seedling shoots and epidermal peels with *HvGEF14* showing the highest fragment counts in these two tissues compared to all other barley *PRONE-GEFs* (Tab. S6).

The leaf epidermis of barley provides an important interface for plant-pathogen interaction. Since the barley type I ROP *Hv*RACB has been shown to play a crucial role in epidermis development and susceptibility to the powdery mildew fungus (Schultheiss *et al*., 2002), we concentrated on PRONE-GEFs which might be of importance to signal transduction in the epidermis. We checked the gene expression of *HvGEF14* via qRT-PCR in leaf and leaf epidermis from seven days old leaves of the barley cultivar Golden Promise. In three independent biological replicates, we found higher transcript levels of *HvGEF14* in the epidermis when compared to whole leaves (Fig. 3A). Notably, in three fully independent biological experiments the *HvGEF14* expression level in the epidermis decreased after inoculation with the biotrophic powdery mildew fungus *Bgh* compared to epidermal peels from unchallenged leaves (Fig. 3B).

**Fig. 3:**
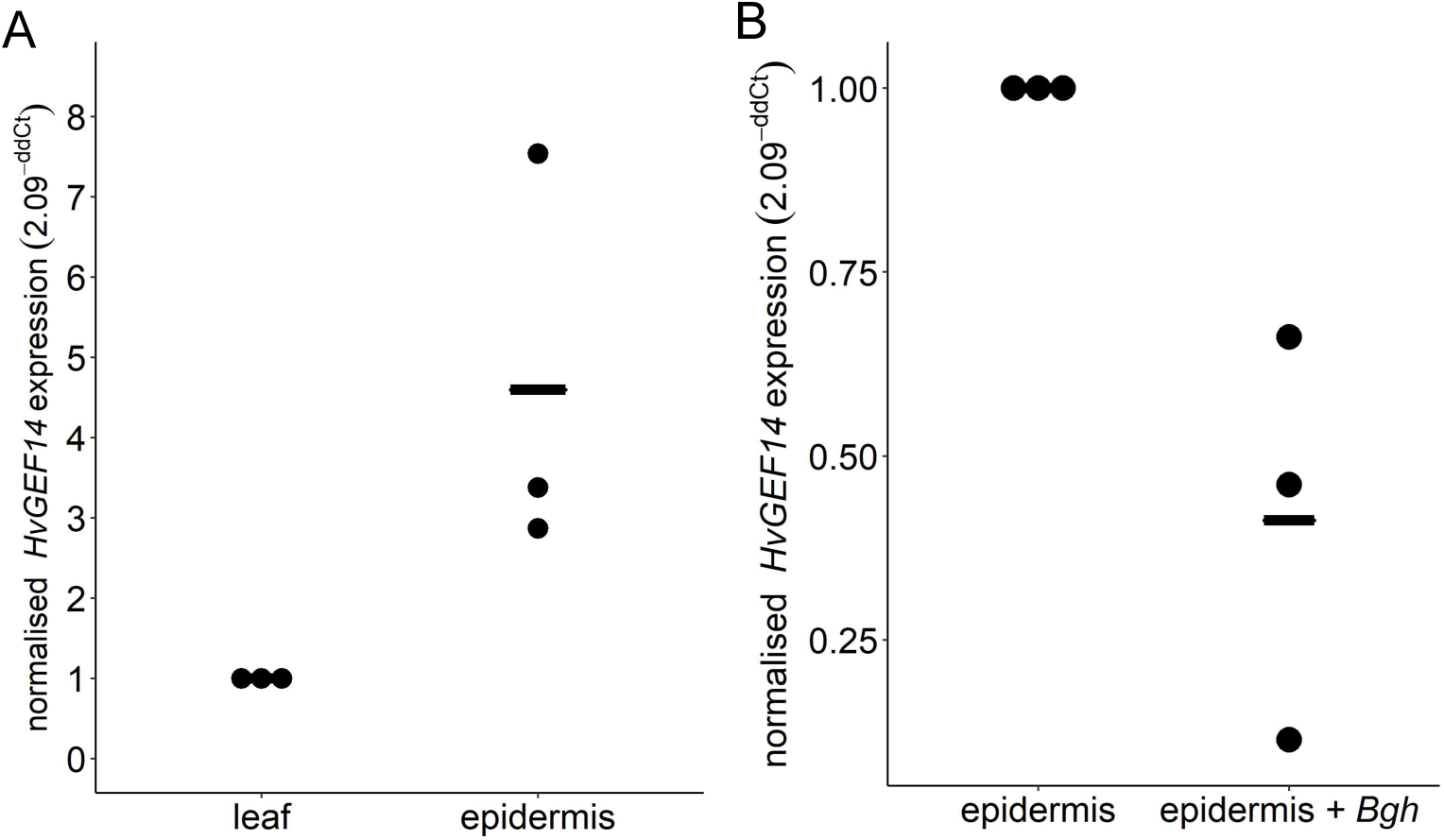
*HvGEF14* shows increased expression in epidermal peels and downregulation after inoculation with fungal spores. *HvGEF14* is expressed in barley epidermis (A) and downregulated after Bgh inoculation (1 dpi) (B) in three independent repetitions (plants harvested on different days). Foldchange expression was calculated with primer efficiency correction and the 2^(-ddCt) method by Livak and Schmittgen (2001) and normalised to transcript levels in whole leaves (A) and non-inoculated epidermis (B).

### *Hv*GEF14 interacts with *Hv*RACB

*Hv*RACB is a barley ROP that supports fungal penetration by *Bgh* into epidermal cells, possibly by supporting polar ingrowth of the fungal haustorium (Engelhardt *et al*., 2020). So far, however, nothing is known about a potential PRONE-GEF-mediated activation of *Hv*RACB. To check if *Hv*GEF14 is the putative HvRACB-activating PRONE-GEF, we analysed the direct protein-protein interaction between *Hv*RACB and *Hv*GEF14 in yeast and *in planta*.

Plant ROPs can be mutagenized in the GTPase domain by exchanging a conserved glycine residue to the hydrophobic valine (e.g., *Hv*RACB-G15V). These mutations render the ROP constitutively activated (CA) regarding downstream signalling because of inactivating the intrinsic GTPase function, which leads to a constantly GTP bound state (Schultheiss *et al*., 2003). Correspondingly, the exchange of a conserved threonine in the N-terminal part of the protein with asparagine keeps the ROP in a dominant negative (DN) confirmation (*Hv*RACB-T20N). A third mutation (*Hv*RACB-D121N) can be performed at an aspartic acid to an uncharged asparagine in order to lower nucleotide affinity and potentially increase GEF-binding affinity (Akamatsu *et al*., 2013; Berken *et al*., 2005; Cool *et al*., 1999).

Interestingly, we found that full length *Hv*GEF14 and the *Hv*GEF14 PRONE domain (amino acids 124-485) directly interact in yeast with wild type *Hv*RACB, CA *Hv*RACB-G15V and the low nucleotide affinity version *Hv*RACB-D121N, but not with the DN *Hv*RACB-T20N variant (Fig. 4). All *Hv*RACB variants were truncated at the *Hv*RACB CSIL motif to inhibit prenylation and membrane association in yeast and hence facilitate protein accumulation in yeast nuclei. To substantiate the results, we performed Western Blots, which confirmed protein stability in yeast (Fig. S5).

**Fig. 4:**
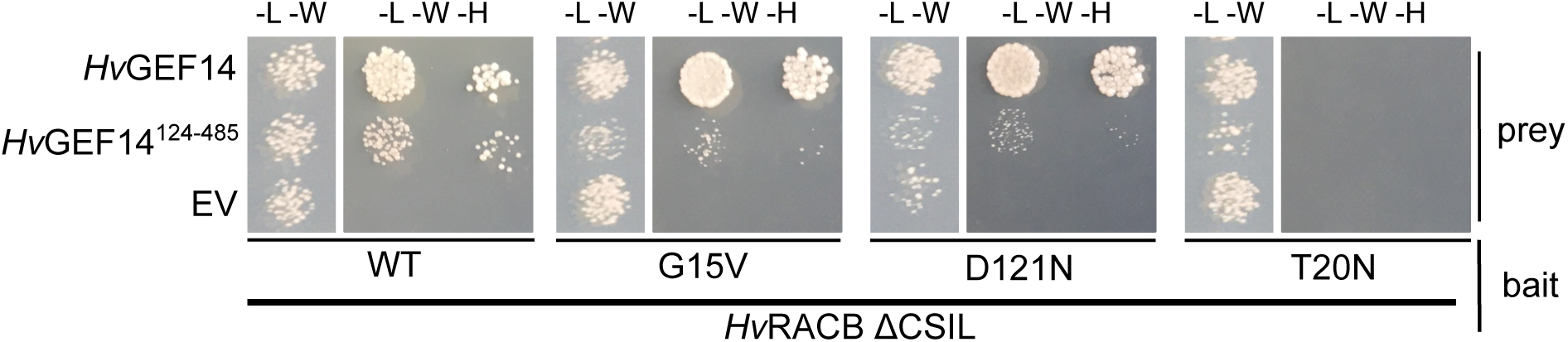
HvGEF14 interacts with barley ROP variants in yeast. *Hv*RACB wild type (WT), G15V constitutively activated (CA), T20N dominant negative (DN) and D121N low nucleotide affinity variants were tested as bait against prey constructs *Hv*GEF14 full length, *Hv*GEF14 amino acids 124-485 (PRONE domain) or empty vector (EV). Interaction of proteins shown on media containing amino acid mix without leucine (L), tryptophan (W) and histidine (H), (-L-W-H) in two dilutions (factor 10^-1^) to identify growth of single yeast colonies. Representative image of three experiments with the same result. Successful yeast transformation was confirmed with selective media (amino acid mix without leucine (L) and tryptophan (W) (-L-W). Dropout was performed on one -L-W and one -L-W-H plate and images were cropped during figure preparation for better visibility. Original images can be found in Fig. S1 (Fig. S4).

To test direct protein-protein interaction between full-length *Hv*GEF14 and previously used variants of *Hv*RACB *in planta*, we conducted a series of experiments measuring Förster resonance energy transfer by fluorescence lifetime imaging (FRET-FLIM) in transiently transformed barley epidermal cells. (Fig. 5). As a FRET fluorophore pair, monomeric enhanced green fluorescent protein (meGFP) and red fluorescent protein mCherry were chosen for their proven effectiveness in plant protein-protein interaction studies (Denay *et al*., 2019). meGFP-*Hv*GEF14 served as FRET-donor and hence was used unchanged as an interaction partner in every combination tested. As FRET acceptor, different variants of mCherry-*Hv*RACB (WT, G15V, D121N, T20N) and free mCherry were used. All measurements took place at the cell periphery at the equatorial cell plane and both fluorophores were detected for the measurements in every case.

**Fig. 5:**
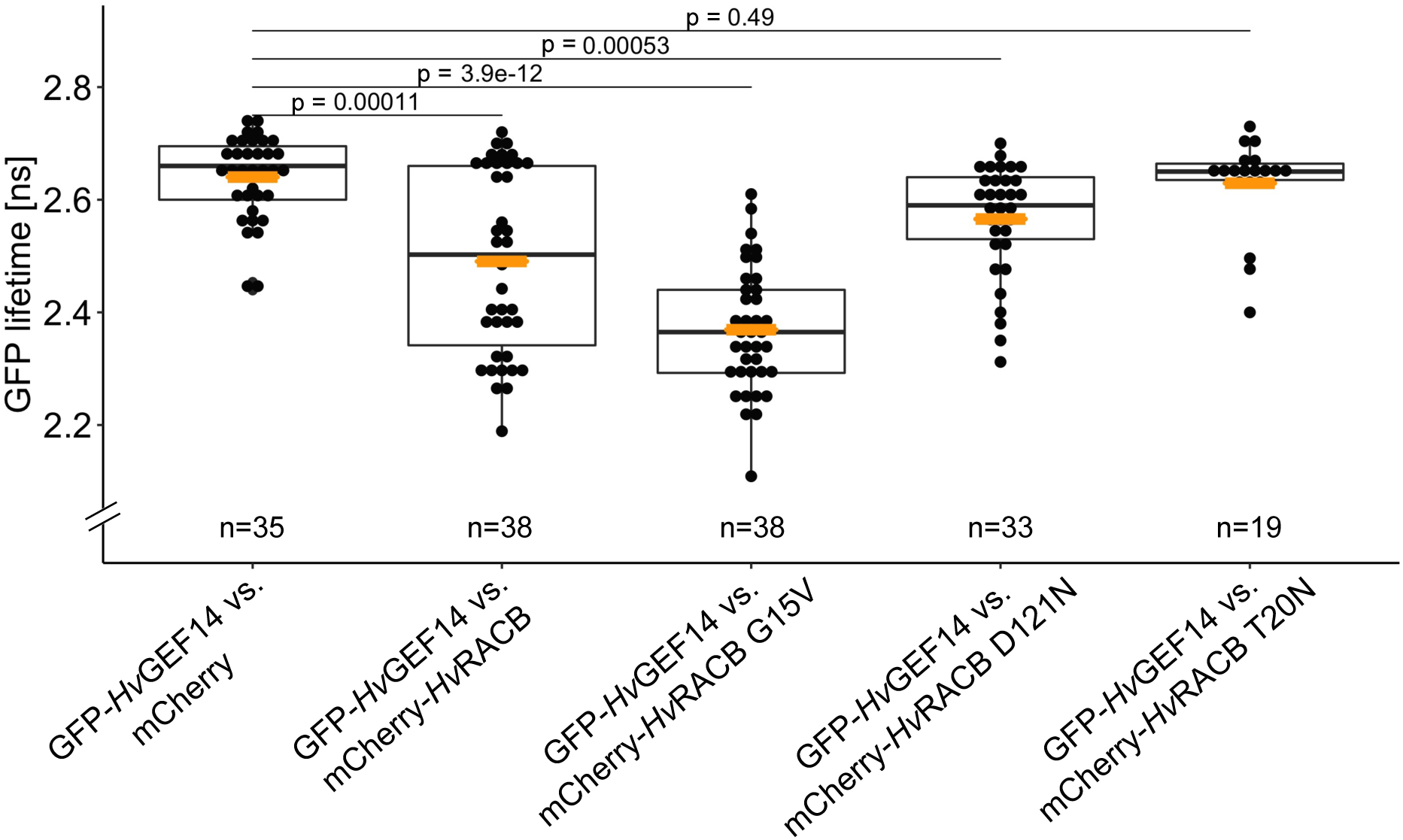
*Hv*GEF14 interacts with HvRACB variants in planta. GFP-tagged *Hv*GEF14 full length was observed in FRET-FLIM experiments to interact with mCherry-tagged *Hv*RACB wild type (WT), constitutively activated (G15V) and low nucleotide affinity mutant (D121N) but not with dominant negative (T20N) RACB. Mean GFP lifetime is indicated with black point, n = number of cells observed in three independent experiments. Mean shown as orange dot. Kruskal-Wallis p-value = 4.346e-14, pairwise comparison was performed via wilcoxon test and bonferroni adjustment for multiple testing. (Rstudio, version 1.2.5033).

Using these donor/acceptor combinations, decreased meGFP-*Hv*GEF14 lifetime can be tested upon co-expression with a particular mCherry-*Hv*RACB variant. In three repetitions, meGFP-*Hv*GEF14 lifetimes varied but showed a significant lifetime reduction in epidermal cells transiently co-expressing meGFP-*Hv*GEF14 and mCherry-*Hv*RACB WT to 2.49 ns on average compared to the negative control with free mCherry recorded at 2.64 ns. This suggests that *Hv*GEF14 interacts directly with wild type *Hv*RACB *in planta* even though the fluorescence lifetime reduction varied in between cells (Fig. 5). The transient co-expression of meGFP-*Hv*GEF14 with mCherry-CA *Hv*RACB-G15V also resulted in a significantly reduced GFP-lifetime of 2.37 ns on average. This reflects the interaction assays in yeast and provides further evidence that *Hv*GEF14 can also interact with the CA variant *Hv*RACB-G15V *in planta* (Fig. 5). Similarly, co-expression of mCherry-fused *Hv*RACB-D121N mutant significantly decreases the meGFP-*Hv*GEF14 lifetime to 2.57 ns on average. However, this combination of proteins leads to a higher population of cells with meGFP-*Hv*GEF14 lifetimes similar to control levels, demonstrating non-interacting proteins. However, these results confirmed the direct interaction of *Hv*GEF14 with *Hv*RACB-D121N in yeast (Fig. 4). The only mCherry-tagged *Hv*RACB variant, which did not provoke a reduced meGFP-*Hv*GEF14 lifetime when compared to the control, was the DN *Hv*RACB-T20N mutant. As observed in yeast-two-hybrid assays, there was no measurable interaction between *Hv*GEF14 and *Hv*RACB-T20N *in planta* (Fig. 5). In total, the FRET-FLIM experiments suggest a direct interaction between *Hv*GEF14 full length and *Hv*RACB wild type, -G15V and -D121N *in planta*.

To test for possible specificity of *Hv*GEF14-ROP interaction, we also tested the interaction with *Hv*RAC1, another epidermis-expressed barley ROP that is of type II. In Y2H, *Hv*GEF14 interacted with WT and the CA *Hv*RAC1-G23V but not with DN *Hv*RAC1-T28N (Fig. S6, S7).

### *Hv*GEF14 over expression leads to activation of barley ROPs

Several ROP interacting proteins display a change of subcellular localisation in presence of an activated ROP. This change in localisation is considered an indirect evidence for local ROP activity, because those interactors preferable interact with GTP-loaded, signalling-activated ROPs (Li *et al*., 2020; McCollum *et al*., 2020; Schultheiss *et al*., 2008). In transiently transformed epidermal cells, CA *Hv*RACB is partially located at the cell periphery/plasma membrane, which depends on its C-terminal CAAL prenylation motif (Schultheiss *et al*., 2003; Schultheiss *et al*., 2008). *Hv*RIC171, a barley scaffold protein that has been shown to directly interact with CA but not DN barley ROP variants (Schultheiss *et al*., 2008), is recruited from the cytoplasm to the cell periphery and plasma membrane in the presence of co-expressed CA *Hv*RACB-G15V but not DN *Hv*RACB-T20N. This strongly suggests *Hv*RACB activation-dependent recruitment of *Hv*RIC171 (Schultheiss *et al*., 2008). Based on this, we monitored the localization of mCherry-tagged *Hv*RIC171 in barley epidermis cells in the presence or absence of co-expressed *Hv*GEF14 to indirectly analyse the potential of *Hv*GEF14 to activate barley ROPs like *Hv*RACB. mCherry-*Hv*RIC171 fluorescence significantly increased at the cell periphery when *Hv*GEF14 was present compared to the empty vector (EV) control (Fig. 6A and B). This was evident from an increase in normalised fluorescent signal intensity at the cell periphery in the equatorial plane of the cell. Additionally, mCherry-*Hv*RIC171 signals appeared very irregular in the cell periphery of control cells whereas mCherry-*Hv*RIC171 more evenly labelled the cell periphery of cells co-expressing the putative ROP activator *Hv*GEF14 (Fig. 6A, lower panel).

**Fig. 6:**
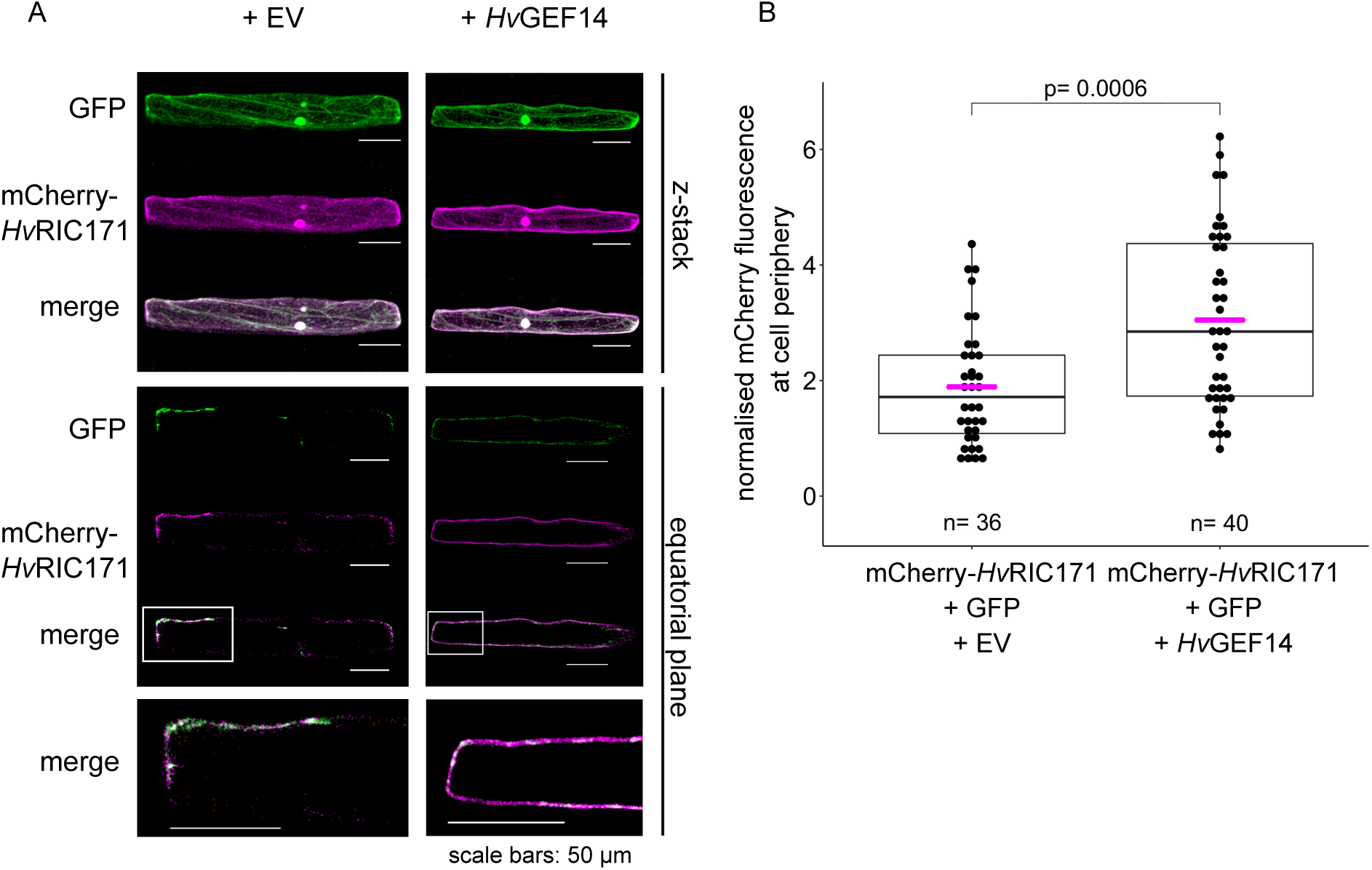
*Hv*GEF14 influences subcellular localisation of mCherry-*Hv*RIC171. Localisation of mCherry-*Hv*RIC171, a downstream interactor of activated ROPs in barley, shows higher fluorescence intensity at the cell periphery when *Hv*GEF14 is co-expressed simultaneously. (A) Representative images of three biological replicates show z-stack and equatorial plane of mCherry-*Hv*RIC171 and cytosolic GFP fluorescence of transiently transformed barley epidermis cells. (B) Quantification of periphery mCherry fluorescence intensity normalized to whole cell mCherry and GFP fluorescence intensity measured in minimum 30 cells via Fiji, statistical analysis performed in Rstudio via wilcox test (Rstudio *version 1.2.5033*).

To test specific activation of the susceptibility factor *Hv*RACB *in planta*, we made use of a FRET-based activity sensor probe containing N-terminal meGFP-tagged *Hv*RACB and C-terminal mCherry-tagged *Hv*CRIB46 on individual plasmids. Barley *Hv*CRIB46 represents an extended CRIB domain of HvRIC171 and was shown before to interact with CA *Hv*RACB but not DN *Hv*RACB. It can hence be considered, such as CRIB domains in general, as preferably interacting with GTP-loaded RACB. To reduce interference of endogenous signalling components in barley, the measurements were performed in *N. benthamiana. meGFP-Hv*RACB WT did not interact with the negative control GST-mCherry (GFP lifetime of 2.62 ns on average), but meGFP-*Hv*RACB WT lifetime is slightly but significantly reduced when co-expressed with *Hv*CRIB46-mCherry (2.57 ns on average). This likely reflects the fact that wild type HvRACB can switch between GDP and GTP loaded forms also in *N. benthamiana*. However, when co-expressing HA-tagged *Hv*GEF14, we measured a significant decrease in meGFP fluorescence lifetime, suggesting abundance of activated CRIB-binding *Hv*RACB-GTP. Since HA-*Hv*GEF14 transformation could not be verified in individual cells via microscopy, the protein was detected via Western Blot after FRET-FLIM measurements (Fig. S8). As a positive control, the interaction of meGFP-*Hv*RACB CA with *Hv*CRIB46-mCherry was measured at 2.35 ns on average (Fig. 7).

**Fig. 7:**
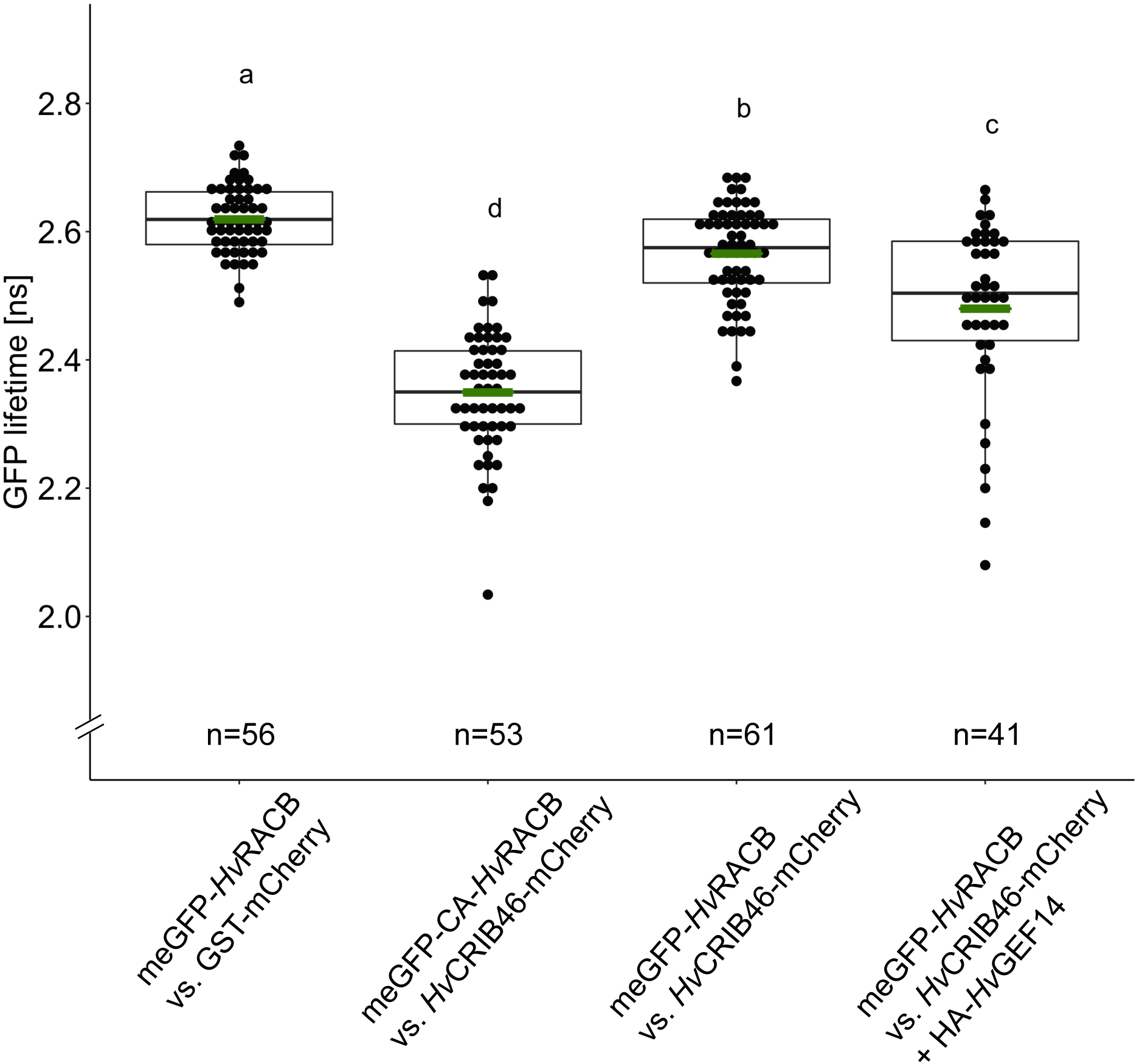
*Hv*RACB can be activated by *Hv*GEF14. meGFP-*Hv*RACB interacts with *Hv*CRIB46-mCherry, a 46 amino acid fragment of *Hv*RIC171, when mutated (G15V) to a constitutively activated variant or in presence of *Hv*GEF14 during FRET-FLIM measurements in *N. benthamiana*. GST-mCherry used as negative control. Summary of six independent repetitions, Kruskal-Wallis comparison of means performed in Rstudio (version 1.2.5033) after testing for distribution of data.

### *Hv*GEF14 supports barley susceptibility towards penetration by *Bgh*

Since *HvGEF14* is expressed in barley shoot epidermis and interacts with two epidermis expressed *Hv*ROPs, we tested the function of the exchange factor in the interaction of barley and *Bgh*. Via transient single cell overexpression or RNAi-mediated silencing of *Hv*GEF14 in barley epidermis cells and subsequent inoculation with fungal spores, we scored interaction outcome for each transformed and attacked cell in at least five independent experiments. On average we observed a significant increase of fungal success from 34.6% to 46.5% penetration efficiency observed as haustorium formation in *Hv*GEF14 overexpressing cells when compared to an empty vector control (Fig. 8).

**Fig. 8:**
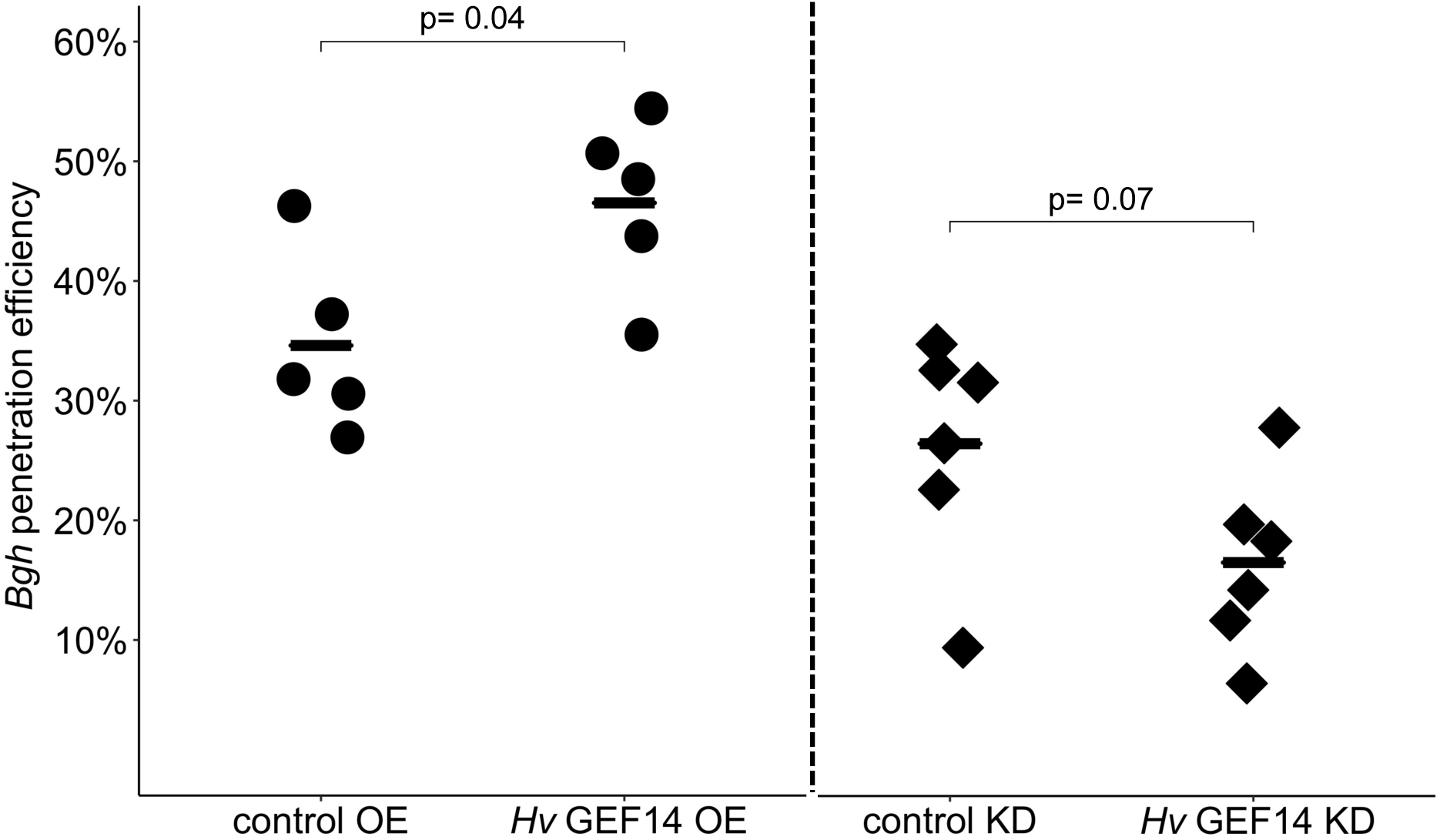
*Hv*GEF14 affects *Bgh* penetration efficiency. *Bgh* penetration efficiency in barley epidermis cells over-expressing (OE) *Hv*GEF14 full length compared to mean penetration efficiency in cells with pGY1 empty vector (control OE) or after knock-down (KD) of *HvGEF14* via RNAi compared to mean penetration efficiency of pIPKTA30N empty vector (control KD). Mean values indicated with bar. Statistical significance of differences of the mean calculated with t-test in Rstudio (version 1.2.5033) after assessing for normal distribution with Shapiro-Wilk test.

We assessed the efficacy of RNAi-mediated gene silencing to 53% according to measuring the reduction of GFP fluorescence intensities of single GFP-*Hv*GEF14 expressing barley cells co-transformed with empty vector or *HvGEF14* gene silencing construct, respectively (Tab. S7). The knock down of *HvGEF14* by RNAi led to the opposite effect of over-expression; a decrease in penetration rate from 26.4% in the controls to only 16.5% in cells, in which *HvGEF14* was silenced by RNAi. Absolute penetration rates are a bit lower in RNAi experiments due to longer incubation time between microprojectile-mediated transformation and inoculation and considerably varied from experiment to experiment even in the controls. Accordingly, a p-value of 0.07 was computed during statistical analysis of the means (Fig. 8). However, when comparing to individual controls, all six independent repetitions of the experiment showed the same trend towards reduced penetration success after knock-down of *HvGEF14*.

## Discussion

The barley susceptibility factor *Hv*RACB has been studied over the last decades to understand underlying molecular mechanisms of its role in support of fungal entry into barley epidermal cells (Engelhardt *et al*., 2020). However, a critical step in this pathway may be the transition from a GDP-bound, signalling inactive, towards the GTP-loaded signalling activated state of *Hv*RACB. As shown in the model species *A. thaliana* and in *O. sativa*, it is apparent that plant specific PRONE-GEFs can facilitate the activation of ROP downstream signalling. In this work, we therefore investigated the role of a barley PRONE-GEF candidate. This revealed the epidermis-expressed and transcriptionally *Bgh*-regulated *Hv*GEF14 and its likely role in ROP activation and susceptibility to fungal penetration.

According to our phylogenetic analysis, we suggest that PRONE-GEF14 has a unique position in the evolution of PRONE-GEFs. A BLAST search with the primary sequence of *At*GEF14 reveals that the closest ortholog to PRONE-GEF14 in the moss *Physcomitrium patens* is *Pp*GEF1 and that there is no PRONE-GEF14 found in the moss (see Eklund *et al*. (2010)). In addition, in the liverwort *Marchantia polymorpha*, only one PRONE-GEF can be found in the genome. The primary sequence of this PRONE-GEF, KARAPPO, is also most similar to *At*GEF1 (Fig. S8 and Hiwatashi *et al*. (2019). However, the ancient angiosperm species *Amborella* seems to encode a GEF14 ortholog (protein accession XP_006878646). Hence, PRONE-GEF14 proteins might have evolved early in angiosperms before separation of monocots and dicots. We could show the abundance of *HvGEF14* transcript in barley leaves and measured higher transcript levels in epidermal peels when compared to whole leaves (Fig. 3A). Possible candidate ROPs to interact with the gene product of *HvGEF14* in barley epidermis are the five epidermis expressed small GTPases: *HvRACB*, *HvRAC1*, *HvRAC3*, *HvRACD* and *HvROP6*. Furthermore, *HvRACB*, *HvRACD* and *HvROP6* had stronger transcript signals in the epidermis compared to mesophyll in RT-PCR assays. When leaves were challenged with *Bgh*, several susceptibility related barley ROP genes had slightly lower transcript levels compared to non-inoculated controls (Schultheiss *et al*., 2003). This gene expression profile in the epidermis is reminiscent of the *HvGEF14* expression in barley epidermis during *Bgh* attack (Fig. 3B).

In the model plant *A. thaliana*, *AtGEF14* expression has also been reported in leaves via RT-PCR experiments. In addition, *AtGEF14* expression patterns responded to a variety of abiotic stresses (Shin *et al*., 2009) which points to underlying regulatory mechanisms in response to environmental signals. It is shown that a *Bgh* effector, *Bgh* ROP-interactive peptide 1, can interact with *Hv*RACB and supports fungal virulence (Nottensteiner *et al*., 2018). We speculate that downregulation of *HvGEF14* transcripts and some barley *ROPs* could reflect a plant response to fungal interference with host ROP signalling, to which the plant reacts by countermeasures and downregulation of the susceptibility pathway. Alternatively, downregulation of *HvGEF14* could reflect a re-allocation of resources from growth into defence, if *Hv*GEF14 would have a physiological function in growth and development.

This study shows direct protein-protein interaction of *Hv*GEF14 with wild type *Hv*RACB, CA *Hv*RACB-G15V and *Hv*RACB-D121N (low nucleotide affinity) (Fig. 4 and Fig. 5). In addition, we report *Hv*GEF14 interaction with wild type *Hv*RAC1 and CA *Hv*RAC1-G23V in yeast (Fig. S6). The fact that *Hv*GEF14 can interact with both, type I and type II ROPs suggests that it might function in more than one signalling pathway. The interaction with WT ROPs provides the basis for a possible role of this barley PRONE-GEF in small GTPase signalling.

In accordance with other studies in *A. thaliana*, which have previously shown interaction of PRONE-GEFs with D121N-like low nucleotide affinity mutants of *At*ROPs (Akamatsu *et al*., 2013; Berken *et al*., 2005; Denninger *et al*., 2019; Gu *et al*., 2006), we could measure *in planta* interaction of *Hv*RACB D121N with *Hv*GEF14. The initial discovery of the 14 *A. thaliana* PRONE-GEFs was made in a yeast-two-hybrid screen with *At*ROP4 D121N as bait (Berken *et al*., 2005). A similar strategy was used to find PRONE-GEFs of *Os*RAC1 in *O. sativa* (Akamatsu *et al*., 2013). Furthermore, a global investigation into *At*GEF-*At*ROP1 interactions showed that *At*GEF14 preferably interacts with a D121A/C188S mutant of *At*ROP1 *in vitro* (Gu *et al*., 2006). This interaction of PRONE-GEFs with low nucleotide affinity ROPs might stem from the fact that these ROP mutants might still cycle between activated and inactivated conformations while providing stronger affinity to PRONE-GEFs (Berken *et al*., 2005). Alternatively, the D121N mutation might lead to a similar protein conformation as the nucleotide free ROP, which has been crystallised in complex with the PRONE domain of *At*GEF8 (Thomas *et al*., 2007).

The interaction of PRONE-GEFs with other mutants of ROPs was also reported in *O. sativa*. *Os*GEF1 interaction with *Os*RAC1 mutants was shown via split Venus fluorescence complementation assays in protoplasts. Both, the constitutively activated *Os*RAC1-G19V as well as the dominant negative *Os*RAC1-T24N variants were able to reconstitute fluorescence of fluorescent Venus at the plasma membrane (Akamatsu *et al*., 2013).

Interaction of PRONE-GEFs with CA ROPs has also been repeatedly reported here and before (e.g., Akamatsu et al. 2013; Denninger et al. 2019). Together, data support that *Hv*GEF14 can interact with *Hv*RACB in *planta*. Together with data from the literature, however, a picture emerges that supports that PRONE-GEFs do not only interact with ROP-GDP but possibly can stay in contact to ROP-GTP after nucleotide exchange. This could have several reasons, which might be further studied. One possibility is that a PRONE-GEF-ROP(GTP) complex is stabilised when the ROP is arrested in a CA status. Alternatively, activated ROPs might stay in contact to GEFs for a feedback regulation on ROP activity through complexes formed with ROP executers as discussed before (Wu and Lew, 2013). Because PRONE-GEFs including *Hv*GEF14 can form dimers, GEF-GEF – ROP-GTP interaction could also recruit further ROP-GDP for activation (Smokvarska *et al*., 2021).

Sub-cellular localisation of signalling protein complexes is a key to ROP-mediated processes. After activation, ROPs relocate to or are stabilized in their association with the plasma membrane. There, they interact with downstream effectors/executers to facilitate, amongst other functions, polar growth processes (Kawano *et al*., 2014; Poraty-Gavra *et al*., 2013; Schultheiss *et al*., 2003). During *A. thaliana* pollen tube growth, for example, the localisation of activated *At*ROP1 to the apical plasma membrane regulates tip growth (Gu *et al*., 2004). This specific localisation of activated *At*ROP1 to the membrane leads to the accumulation of downstream executers like *At*RIC4 in the same compartment. In presence of dominant negative *At*ROP1, however, the RIC protein localises to the cytoplasm (Gu *et al*., 2005). In addition, the co-expression of a ROP-GAP, which renders *At*ROP1 inactive, also results in the relocation of *At*RIC4 to the cytoplasm. In this model system of polar growth, the correct localisation of activated *At*ROP1 as well as *At*RIC4 was observed to be crucial for downstream signal transduction and fine tuning of the growth process (Hwang *et al*., 2005). To assess *Hv*RACB signalling activity status *in planta*, we made use of the previously published recruitment of *Hv*RIC171 to CA *Hv*RACB-G15V, which otherwise largely resides at the plasma membrane. In the same studies, *Hv*RIC171 was shown to be recruited to the site of fungal penetration likely via endogenously activated ROPs (Schultheiss *et al*., 2003; Schultheiss *et al*., 2008). Via confocal microscopy, we measured higher mCherry-*Hv*RIC171 localisation at the cell periphery in the presence of transiently over-expressed *Hv*GEF14 (Fig. 6). Based on previous work on *A. thaliana* and *H. vulgare* RICs, this finding suggests that the over-expression of *Hv*GEF14 leads to a higher ratio of activated endogenous ROPs, which in turn recruit *Hv*RIC171 towards the cell periphery. Another way to observe ROP activation *in planta* is via CRIB-based ROP activity sensors that have previously been adapted for plants (Kawano *et al*., 2010; Wang *et al*., 2018; Wong *et al*., 2018). This so-called Ras and interacting chimeric unit (Raichu) sensor includes the fluorophores Venus and CFP as FRET-pair. ROP activity is monitored by its interaction with the CRIB domain of a downstream executer. In a similar manner, we used the interaction of activated *Hv*RACB with a *Hv*RIC171 fragment containing the CRIB domain in FRET-FLIM assays using the efficient FRET-pair meGFP and mCherry on separate plasmids (Denay *et al*., 2019). With the setup in this study, it is possible to monitor ROP activation status in *N. benthamiana* cells with different ROP and interactor combinations. We observed a lower GFP-lifetime when we co-expressed *Hv*GEF14, which strongly suggests that it can activate *Hv*RACB which leads to interaction of GTP-loaded *Hv*RACB and *Hv*CRIB46 (Fig. 7).

In parallel, the transient over-expression of *Hv*GEF14 in barley epidermal cells leads to a significant increase in *Bgh* penetration success (Fig. 8). This phenotype is similar to enhanced fungal penetration during CA *Hv*RACB-G15V over-expression (Schultheiss *et al*., 2003), as well as the over-expression of *Hv*RACB-downstream executers, such as *Hv*RIC171 (Schultheiss *et al*., 2008) and *Hv*RIPb (McCollum *et al*., 2020). Transient silencing of *Hv*GEF14, on the other hand, renders barley epidermis cells by trend more resistant to *Bgh* penetration (Fig. 8). Even though the phenotype could be observed in every repetition when comparing *Hv*GEF14 RNAi with its respective control, the extend of fungal penetration varied amongst repetitions so that the significance threshold of 0.05 was not met after statistical analysis. Considering however, that the inoculation with *Bgh* naturally decreases the transcript of *HvGEF14* (see Fig. 3), additional ectopic knock down perhaps cannot be expected to exert a major additional effect. Taken together, these assays point to a role of *Hv*GEF14 in supporting the accommodation of *Bgh* infection structures in barley epidermal cells, similar to and possibly in cooperation with the susceptibility factor *Hv*RACB and other epidermis expressed *Hv*ROPs (Schultheiss *et al*., 2003).

An overview and comprehensive annotation of the PRONE-GEFs from *A. thaliana*, *O. sativa* and *H. vulgare* was prepared using maximum likelihood phylogenetic analysis. The analysed PRONE-GEFs distribute into three clades based on high bootstrap values at the branching points (Fig. 1) which is partly in accordance with previously published PRONE-GEF phylogenies (Berken *et al*., 2005; Kim *et al*., 2020; Shin *et al*., 2009; Zhang and McCormick, 2007). Shin and colleagues (2009) use the same alignment algorithm as in this study to build a dendrogram of *A. thaliana* PRONE-GEFs. The resulting tree does not include bootstrap analysis but also highlights three clades of PRONE-GEFs with different distribution when compared to the clades published in this study. In their analysis, *At*GEF14 belongs to group 2, and *At*GEF1 is most separated from all other *A. thaliana* PRONE-GEFs (Shin *et al*., 2009). Our results, however, indicate that PRONE-GEF1 from all three species cluster in clade I (Fig. 1). *At*GEF14 is placed next to *At*GEF12 and *At*GEF11 in Shin et al. (2009). This is also the case in Berken *et al*. (2005) who report a phylogram of *A. thaliana*, *O. sativa*, *M. truncatula* (*Mt*) and *L. esculentum* (*Le*) PRONE-GEFs. The resulting TreeView phylogeny in Berken *et al*. (2005) also does not report bootstrap values and differs to our phylogenetic tree in the early branching of *At*GEF7. Another phylogeny has been published by Zhang and McCormick (2007) who compare to that date known PRONE-GEFs from *A. thaliana*, *O. sativa*, *P. patens* (*Pp*), *M. truncatula* (*Mt*) and *L. esculentum* (*Le*). Based on CLUSTALW alignment and MEGA3.1 neighbour joining algorithm with bootstrap analysis, the authors define two distinct groups of PRONE-GEFs with *At*GEFs 8, 9, 11, 12, 13 and 14 as well as *Le*KPP and *Mt*GEF1 in one group with several non-annotated *Os*GEFs. In addition to the two groups of PRONE-GEFs published by Zhang and McCormick (2007), we emphasize the branching of PRONE-GEF14 proteins (with high confidence of bootstrap value 100) by dividing previously published clade II further and highlighting the unique position of PRONE-GEF14 in a separate clade III. A more recently published PRONE-GEF alignment compares *A. thaliana* and *O. sativa* proteins (Kim *et al*., 2020) with similar methods as has been used here. Though the annotation of *Os*GEFs does not match with previously published *Os*GEF nomenclature we can confirm distinct clades of PRONE-GEFs 1-7 and 8-13 as well as the placement of PRONE-GEF14 separated from the other two clades (Fig. 1 and (Kim *et al*., 2020)). Due to high confidence bootstrap analysis and comprehensive annotation of *O. sativa* and *H. vulgare* PRONE-GEFs, we propose to base further comparisons on the presented phylogeny (Fig. 1). A unique position of GEF14 proteins in the phylogeny of PRONE-GEFs is further supported by its comparably low sequence conservation of the PRONE domain, when compared to all other PRONE-GEFs (Fig. S1, Tab. S1).

In conclusion, *Hv*GEF14 is a *bona fide* barley PRONE-GEF and interacts with barley ROPs. The interaction with susceptibility-related barley ROPs might lead to ROP activation and therefore facilitate ROP functions in susceptibility to invasion by *Bgh*. Research on *A. thaliana* PRONE-GEFs highlights the interplay of different PRONE-GEFs to facilitate polar growth processes like root hair formation or pollen tube growth (Denninger *et al*., 2019; Li *et al*., 2020). It has to be studied if *Hv*GEF14 function is choreographed in a similar manner with other barley PRONE-GEFs. In addition, other PRONE-GEF interactors, such as potential upstream RLKs, remain to be investigated. In future studies, susceptibility- and *Hv*RACB-related candidate RLKs (Douchkov *et al*., 2014; Schnepf *et al*., 2018) will be of special interest to link *Hv*RACB signalling to cell surface signal perception.

## Supporting information

Full supplementary data

## Supplementary data

Fig. S1: Phylogenetic tree of *A. thaliana* and *O. sativa* PRONE-GEFs.

Fig. S2: MUSCLE alignment of *A. thaliana*, *O. sativa* and *H. vulgare* PRONE-GEFs.

Fig. S3: *Hv*GEF14 homo-dimerises with full length and PRONE domain (amino acids 124-485) in yeast-two hybrid.

Fig. S4: Original images of *Hv*GEF14 full length and PRONE domain (amino acids 124-485) interaction with *Hv*RACB variants (wild type, WT; G15V; D121N; T20N) via yeast-two hybrid.

Fig. S5: Western Blot of yeast proteins of Y2H *Hv*GEF14 interaction with *Hv*RACB on PVDF membranes.

Fig. S6: *Hv*RAC1 wild type (WT), G23V constitutively activated (CA), T28N dominant negative (DN) variants were tested as bait against prey constructs *Hv*GEF14 full length, *Hv*GEF14 124-485 (PRONE domain) or empty vector (EV).

Fig. S7: Western Blot of yeast proteins Y2H *Hv*GEF14 interaction with *Hv*RAC1 on PVDF membranes.

Fig. S8: Western Blot of proteins extracted from *N. benthamiana* on PVDF membranes.

Table S1: Primary sequence similarity of HvGEFs when compared to consensus sequence shows highest variability of HvGEF14 primary sequence.

Table S2: List of barley PRONE-GEFs with gene/protein identifiers, annotation, PRONE prediction, and protein sequence length.

Table S3: List of plasmids used in this study.

Table S4: List of primers used in this study.

Table S6: Tissue-specific *Hv*GEF gene expression based on RNAseq (James Hutton Institute) in fragments per reads per kilobase million (FPKM).

Table S7: RNAi efficiency of GEF14 hairpin construct in transient transformation of barley epidermal cells.

## Abbreviations

ROP: Rho of plants
GEF: Guanine nucleotide exchange factor
PRONE: plant specific ROP-nucleotide exchanger
CA: constitutively activated
DN: dominant negative
GAP: GTPase activating protein
RIC: ROP interacting and CRIB domain containing protein
RIP: ROP interactive partner
GDI: Guanine nucleotide dissociation inhibitor
Bgh: Blumeria graminis forma specialis hordei
DH-PH: diffuse B-cell lymphoma homology – pleckstrin homology
DHR2: DOCK homology region 2
RLK: receptor like kinase
FER: FERONIA
Hv: Hordeum vulgare
At: Arabidopsis thaliana
Os: Oryza sativa
Nb: Nicotiana benthamiana
meGFP: monomeric enhanced green fluorescent protein
EV: empty vector

## Acknowledgements

We would like to acknowledge Johanna Hofer for outstanding technical assistance as well as Carolina Galgenmüller. Thanks goes to Parvinderdeep Kahlon and Lina Muñoz for critical reading of the manuscript as well as Lukas Weiss, Christopher McCollum and Michaela Stegmann for intellectual input and methodological assistance. We would like to thank the Center of Advanced Light Microscopy (CALM) imaging centre of TUM for providing access to the FRET-FLIM system. We are grateful to the German Research Foundation (DFG) for funding this work via SFB924.

## Author contributions

RH was responsible for conception and AT, SE and RH designed the work. Data acquisition and analysis was performed by AT as well as interpretation of the data in collaboration with SE and RH. SE assisted in data acquisition for Fig. 8. AT drafted the manuscript and prepared the final version of the work after revision by SE and RH. All authors have approved the submitted version and have agreed both to be personally accountable for the author’s own contributions and to ensure that questions related to the accuracy or integrity of any part of the work, even ones in which the author was not personally involved, are appropriately investigated, resolved, and the resolution documented in the literature.

## Data availability statement

All data generated or analysed during this study are included in this published article [and its supplementary information files]. Additional datasets used and/or analysed during the current study are available from the corresponding author on reasonable request.

## References

1. Akamatsu A, Uno K, Kato M, Wong HL, Shimamoto K, Kawano Y. 2015. New insights into the dimerization of small GTPase Rac/ROP guanine nucleotide exchange factors in rice. Plant signaling and behavior 10, e1044702.

2. Akamatsu A, Wong HL, Fujiwara M, Okuda J, Nishide K, Uno K, Imai K, Umemura K, Kawasaki T, Kawano Y, Shimamoto K. 2013. An OsCEBiP/OsCERK1-OsRacGEF1-OsRac1 module is an essential early component of chitin-induced rice immunity. Cell host & microbe 13, 465–476.

3. Berken A, Thomas C, Wittinghofer A. 2005. A new family of RhoGEFs activates the Rop molecular switch in plants. Nature 436, 1176–1180.

4. Bloch D, Yalovsky S. 2013. Cell polarity signaling. Current opinion in plant biology 16, 734–742.

5. Chang F, Gu Y, Ma H, Yang Z. 2013. AtPRK2 promotes ROP1 activation via RopGEFs in the control of polarized pollen tube growth. Molecular plant 6, 1187–1201.

6. Chen M, Liu H, Kong J, Yang Y, Zhang N, Li R, Yue J, Huang J, Li C, Cheung AY, Tao L-Z. 2011. RopGEF7 Regulates PLETHORA-Dependent Maintenance of the Root Stem Cell Niche in Arabidopsis. The Plant cell 23, 2880–2894.

7. Chen X, Friml J. 2014. Rho-GTPase-regulated vesicle trafficking in plant cell polarity. Biochemical Society transactions 42, 212–218.

8. Chomczynski P, Sachhi N. 1987. Single-step Method of RNA Isolation by Acid Guanidinium Thiocyanate-Phenol-Chloroform Extraction. Analytical Biochemistry 162, 156–159.

9. Cool RH, Schmidt G, Lenzen CU, Prinz H, Vogt D, Wittinghofer A. 1999. The Ras Mutant D119N Is Both Dominant Negative and Activated. MOLECULAR AND CELLULAR BIOLOGY 19, 6297–6305.

10. Denay G, Schultz P, Hansch S, Weidtkamp-Peters S, Simon R. 2019. Over the rainbow: A practical guide for fluorescent protein selection in plant FRET experiments. Plant Direct 3, 1–14.

11. Denninger P, Reichelt A, Schmidt VAF, Mehlhorn DG, Asseck LY, Stanley CE, Keinath NF, Evers JF, Grefen C, Grossmann G. 2019. Distinct RopGEFs Successively Drive Polarization and Outgrowth of Root Hairs. Current Biology 29, 1854–1865 e1855.

12. Douchkov D, Lück S, Johrde A, Nowara D, Himmelbach A, Rajaraman J, Stein N, Sharma R, Kilian B, Schweizer P. 2014. Discovery of genes affecting resistance of barley to adapted and non-adapted powdery mildew fungi. Genome Biology 15.

13. Eklund DM, Svensson EM, Kost B. 2010. Physcomitrella patens: a model to investigate the role of RAC/ROP GTPase signalling in tip growth. Journal of experimental botany 61, 1917–1937.

14. Engelhardt S, Trutzenberg A, Hueckelhoven R. 2020. Regulation and Functions of ROP GTPases in Plant-Microbe Interactions. Cells 9.

15. Fehér A, Lajkó DB. 2015. Signals fly when kinases meet Rho-of-plants (ROP) small G-proteins. Plant Science 237, 93–107.

16. Gu Y, Fu Y, Dowd P, Li S, Vernoud V, Gilroy S, Yang Z. 2005. A Rho family GTPase controls actin dynamics and tip growth via two counteracting downstream pathways in pollen tubes. Journal of Cell Biology 169, 127–138.

17. Gu Y, Li S, Lord EM, Yang Z. 2006. Members of a novel class of Arabidopsis Rho guanine nucleotide exchange factors control Rho GTPase-dependent polar growth. The Plant cell 18, 366–381.

18. Gu Y, Wang Z, Yang Z. 2004. ROP/RAC GTPase: An old new master regulator for plant signaling. Current opinion in plant biology 7, 527–536.

19. Hiwatashi T, Goh H, Yasui Y, Koh LQ, Takami H, Kajikawa M, Kirita H, Kanazawa T, Minamino N, Togawa T, Sato M, Wakazaki M, Yamaguchi K, Shigenobu S, Fukaki H, Mimura T, Toyooka K, Sawa S, Yamato KT, Ueda T, Urano D, Kohchi T, Ishizaki K. 2019. The RopGEF KARAPPO Is Essential for the Initiation of Vegetative Reproduction in Marchantia polymorpha. Current Biology 29, 1–7.

20. Hoefle C, Huesmann C, Schultheiss H, Börnke F, Hensel G, Kumlehn J, Hückelhoven R. 2011. A barley ROP GTPase ACTIVATING PROTEIN associates with microtubules and regulates entry of the barley powdery mildew fungus into leaf epidermal cells. The Plant cell 23, 2422–2439.

21. Hoefle C, McCollum C, Hückelhoven R. 2020. Barley ROP-Interactive Partner-a organizes into RAC1- and MICROTUBULE-ASSOCIATED ROP-GTPASE ACTIVATING PROTEIN 1-dependent membrane domains. BMC Plant Biology 20, 1–12.

22. Huang C, jiao X, Yang L, Zhang M, Dai M, Wang L, Wang K, Bai L, Song C. 2018a. ROP-GEF signal transduction is involved in AtCAP1-regulated root hair growth. Plant Growth Regulation 87, 1–8.

23. Huang J, Liu H, Berberich T, Liu Y, Tao LZ, Liu T. 2018b. Guanine Nucleotide Exchange Factor 7B (RopGEF7B) is involved in floral organ development in Oryza sativa. Rice (N Y) 11, 1–13.

24. Hückelhoven R, Dechert C, Kogel KH. 2003. Overexpression of barley BAX inhibitor 1 induces breakdown of mlo-mediated penetration resistance to Blumeria graminis. Proceedings of National Academy of Science USA 100, 5555–5560.

25. Hwang J-U, Gu Y, Lee Y-J, Yang Z. 2005. Oscillatory ROP GTPase Activation Leads the Oscillatory Polarized Growth of Pollen Tubes. Molecular Biology of the Cell 16, 5385–5399.

26. International Barley Genome Sequencing Consortium T. 2012. A physical, genetic and functional sequence assembly of the barley genome. Nature 491, 711–715.

27. Kawano Y, Akamatsu A, Hayashi K, Housen Y, Okuda J, Yao A, Nakashima A, Takahashi H, Yoshida H, Wong HL, Kawasaki T, Shimamoto K. 2010. Activation of a Rac GTPase by the NLR family disease resistance protein Pit plays a critical role in rice innate immunity. Cell Host Microbe 7, 362–375.

28. Kawano Y, Kaneko-Kawano T, Shimamoto K. 2014. Rho family GTPase-dependent immunity in plants and animals. Frontiers in plant science 5, 522.

29. Kim E-J, Park S-W, Hong W-J, Silva J, Liang W, Zhang D, Jung K-H, Kim Y-J. 2020. Genome-wide analysis of RopGEF gene family to identify genes contributing to pollen tube growth in rice (Oryza sativa). BMC Plant Biology 20, 1.

30. Li E, Zhang YL, Shi X, Li H, Yuan X, Li S, Zhang Y. 2020. A positive feedback circuit for ROP-mediated polar growth. Molecular plant 14, 395–410.

31. Lin W, Tang W, Anderson C, Yang Z. 2018. FERONIA’s sensing of cell wall pectin activates ROP GTPase signaling in Arabidopsis. BioRxiV, 33.

32. Liu J, Liu MX, Qiu LP, Xie F. 2020. SPIKE1 Activates the GTPase ROP6 to Guide the Polarized Growth of Infection Threads in Lotus japonicus. Plant Cell 32, 3774–3791.

33. Livak KJ, Schmittgen TD. 2001. Analysis of relative gene expression data using real-time quantitative PCR and the 2(-Delta Delta C(T)) Method. Methods 25, 402–408.

34. Mascher M, Wicker T, Jenkins J, Plott C, Lux T, Koh CS, Ens J, Gundlach H, Boston LB, Tulpova Z, Holden S, Hernandez-Pinzon I, Scholz U, Mayer KFX, Spannagl M, Pozniak CJ, Sharpe AG, Simkova H, Moscou MJ, Grimwood J, Schmutz J, Stein N. 2021. Long-read sequence assembly: a technical evaluation in barley. Plant Cell 0, 1–19.

35. McCollum C, Engelhardt S, Weiss L, Huckelhoven R. 2020. ROP INTERACTIVE PARTNER b Interacts with RACB and Supports Fungal Penetration into Barley Epidermal Cells. Plant Physiol 184, 823–836.

36. Mergner J, Frejno M, List M, Papacek M, Chen X, Chaudhary A, Samaras P, Richter S, Shikata H, Messerer M, Lang D, Altmann S, Cyprys P, Zolg DP, Mathieson T, Bantscheff M, Hazarika RR, Schmidt T, Dawid C, Dunkel A, Hofmann T, Sprunck S, Falter-Braun P, Johannes F, Mayer KFX, Jurgens G, Wilhelm M, Baumbach J, Grill E, Schneitz K, Schwechheimer C, Kuster B. 2020. Mass-spectrometry-based draft of the Arabidopsis proteome. Nature 579, 409–414.

37. Mucha E, Fricke I, Schaefer A, Wittinghofer A, Berken A. 2011. Rho proteins of plants--functional cycle and regulation of cytoskeletal dynamics. European Journal of Cell Biology 90, 934–943.

38. Nagawa S, Xu T, Yang Z. 2010. RHO GTPase in plants: Conservation and invention of regulators and effectors. Small GTPases 1, 78–88.

39. Nibau C, Wu HM, Cheung AY. 2006. RAC/ROP GTPases: ‘hubs’ for signal integration and diversification in plants. Trends Plant Science 11, 309–315.

40. Nottensteiner M, Zechmann B, McCollum C, Huckelhoven R. 2018. A Barley Powdery Mildew Fungus Non-Autonomous Retrotransposon Encodes a Peptide that Supports Penetration Success on Barley. Journal of experimental botany 69, 293745–293758.

41. Opalski KS, Schultheiss H, Kogel K-H, Hückelhoven R. 2005. The receptor-like MLO protein and the RAC/ROP family G-protein RACB modulate actin reorganization in barley attacked by the biotrophic powdery mildew fungus Blumeria graminis f.sp. hordei. The Plant journal : for cell and molecular biology 41, 291–303.

42. Pathuri IP, Zellerhoff N, Schaffrath U, Hensel G, Kumlehn J, Kogel K-H, Eichmann R, Hückelhoven R. 2008. Constitutively activated barley ROPs modulate epidermal cell size, defense reactions and interactions with fungal leaf pathogens. Plant cell reports 27, 1877–1887.

43. Poraty-Gavra L, Zimmermann P, Haigis S, Bednarek P, Hazak O, Stelmakh OR, Sadot E, Schulze-Lefert P, Gruissem W, Yalovsky S. 2013. The Arabidopsis Rho of plants GTPase AtROP6 functions in developmental and pathogen response pathways. Plant Physiology 161, 1172–1188.

44. Qiu JL, Jilk R, Marks MD, Szymanski DB. 2002. The Arabidopsis SPIKE1 gene is required for normal cell shape control and tissue development. Plant Cell 14, 101–118.

45. Qu S, Zhang X, Song Y, Lin J, Shan X. 2017. THESEUS1 positively modulates plant defense responses against Botrytis cinerea through GUANINE EXCHANGE FACTOR4 signaling. Journal of Integrative Plant Biology 59, 797–804.

46. Scheler B, Schnepf V, Galgenmüller C, Ranf S, Hückelhoven R. 2016. Barley disease susceptibility factor RACB acts in epidermal cell polarity and positioning of the nucleus. Journal of experimental botany 67, 3263–3275.

47. Schmidt A, Hall A. 2002. Guanine nucleotide exchange factors for Rho GTPases: turning on the switch. Genes & Development 16, 1587–1609.

48. Schnepf V, Vlot AC, Kugler K, Hückelhoven R. 2018. Barley susceptibility factor RACB modulates transcript levels of signalling protein genes in compatible interaction with Blumeria graminis f.sp. hordei. Molecular plant pathology 19, 393–404.

49. Schultheiss H, Dechert C, Kogel K-H, Hückelhoven R. 2002. A small GTP-binding host protein is required for entry of powdery mildew fungus into epidermal cells of barley. Plant Physiology 128, 1447–1454.

50. Schultheiss H, Dechert C, Kogel K-H, Hückelhoven R. 2003. Functional analysis of barley RAC/ROP G-protein family members in susceptibility to the powdery mildew fungus. The Plant Journal 36, 589– 601.

51. Schultheiss H, Hensel G, Imani J, Broeders S, Sonnewald U, Kogel K-H, Kumlehn J, Hückelhoven R. 2005. Ectopic expression of constitutively activated RACB in barley enhances susceptibility to powdery mildew and abiotic stress. Plant Physiology 139, 353–362.

52. Schultheiss H, Preuss J, Pircher T, Eichmann R, Hückelhoven R. 2008. Barley RIC171 interacts with RACB in planta and supports entry of the powdery mildew fungus. Cellular microbiology 10, 1815– 1826.

53. Shin DH, Kim T-L, Kwon Y-K, Cho M-H, Yoo J, Jeon J-S, Hahn T-R, Bhoo SH. 2009. Characterization of Arabidopsis RopGEF family genes in response to abiotic stresses. Plant Biotechnology Reports 3, 183–190.

54. Smokvarska M, Jaillais Y, Martiniere A. 2021. Function of membrane domains in Rho-Of-Plant signaling. Plant Physiology 185, 663–681.

55. Tang W, Lin W, Li B, Yang Z. 2021. Mechano-transduction via the pectin-FERONIA complex regulates ROP6 GTPase signaling in Arabidopsis. BioRxiV.

56. Thomas C, Fricke I, Scrima A, Berken A, Wittinghofer A. 2007. Structural evidence for a common intermediate in small G protein-GEF reactions. Molecular cell 25, 141–149.

57. Thomas C, Fricke I, Weyand M, Berken A. 2009. 3D structure of a binary ROP-PRONE complex: The final intermediate for a complete set of molecular snapshots of the RopGEF reaction. Biological chemistry 390, 427–435.

58. Vetter IR, Wittinghofer A. 2001. The guanine nucleotide-binding switch in three dimensions. Science 294, 1299–1304.

59. Wang Q, Li Y, Ishikawa K, Kosami K-I, Uno K, Nagawa S, Tan L, Du J, Shimamoto K, Kawano Y. 2018. Resistance protein Pit interacts with the GEF OsSPK1 to activate OsRac1 and trigger rice immunity. Proceedings of the National Academy of Sciences of the United States of America 115, 1–10.

60. Winge P, Brembu T, Kristensen R, Bones AM. 2000. Genetic Structure and Evolution of RAC-GTPases in Arabidopsis thaliana. Genetics 156, 1959–1971.

61. Wong HL, Akamatsu A, Wang Q, Higuchi M, Matsuda T, Okuda J, Kosami KI, Inada N, Kawasaki T, Kaneko-Kawano T, Nagawa S, Tan L, Kawano Y, Shimamoto K. 2018. In vivo monitoring of plant small GTPase activation using a Forster resonance energy transfer biosensor. Plant Methods 14, 56.

62. Wu C-F, Lew DJ. 2013. Beyond symmetry-breaking: Competition and negative feedback in GTPase regulation. Trends in cell biology 23, 476–483.

63. Wu G, Li H, Yang Z. 2000. Arabidopsis RopGAPs Are a Novel Family of Rho GTPase-Activating Proteins that Require the Cdc42/Rac- Interactive Binding Motif for Rop-Specific GTPase Stimulation. Plant Physiology 124, 1625–1636.

64. Yalovsky S. 2015. Protein lipid modifications and the regulation of ROP GTPase function. Journal of experimental botany 66, 1617–1624.

65. Yamaguchi K, Imai K, Akamatsu A, Mihashi M, Hayashi N, Shimamoto K, Kawasaki T. 2012. SWAP70 functions as a Rac/Rop guanine nucleotide-exchange factor in rice. The Plant Journal 70, 389–397.

66. Yamaguchi K, Kawasaki T. 2012. Function of Arabidopsis SWAP70 GEF in immune response. Plant signaling & behavior 7, 465–468.

67. Yang Y, Li R, Qi M. 2000. In vivo analysis of plant promoters and transcription factors by agroinfiltration of tobacco leaves. The Plant Journal 22, 543–551.

68. Yoo JH, Park J-H, Cho S-H, Yoo S-C, Li J, Zhang H, Kim K-S, Koh H-J, Paek N-C. 2011. The rice bright green leaf (bgl) locus encodes OsRopGEF10, which activates the development of small cuticular papillae on leaf surfaces. Plant molecular biology 77, 631–641.

69. Zhang T, Lei J, Yang H, Xu K, Wang R, Zhang Z. 2011. An improved method for whole protein extraction from yeast Saccharomyces cerevisiae. Yeast 28, 795–798.

70. Zhang Y, McCormick S. 2007. A distinct mechanism regulating a pollen-specific guanine nucleotide exchange factor for the small GTPase Rop in Arabidopsis thaliana. Proceedings of the National Academy of Sciences of the United States of America 104, 18830–18835.

71. Zheng Z-L, Yang Z. 2000. The Rop GTPase: an emerging switch in plants. Plant molecular biology, 1–9.

