## Supplementary material for "Barley guanine nucleotide exchange factor *Hv*GEF14 is an activator of the susceptibility factor *Hv*RACB and supports host cell entry by *Blumeria graminis* f.sp. *hordei*": Full supplementary data

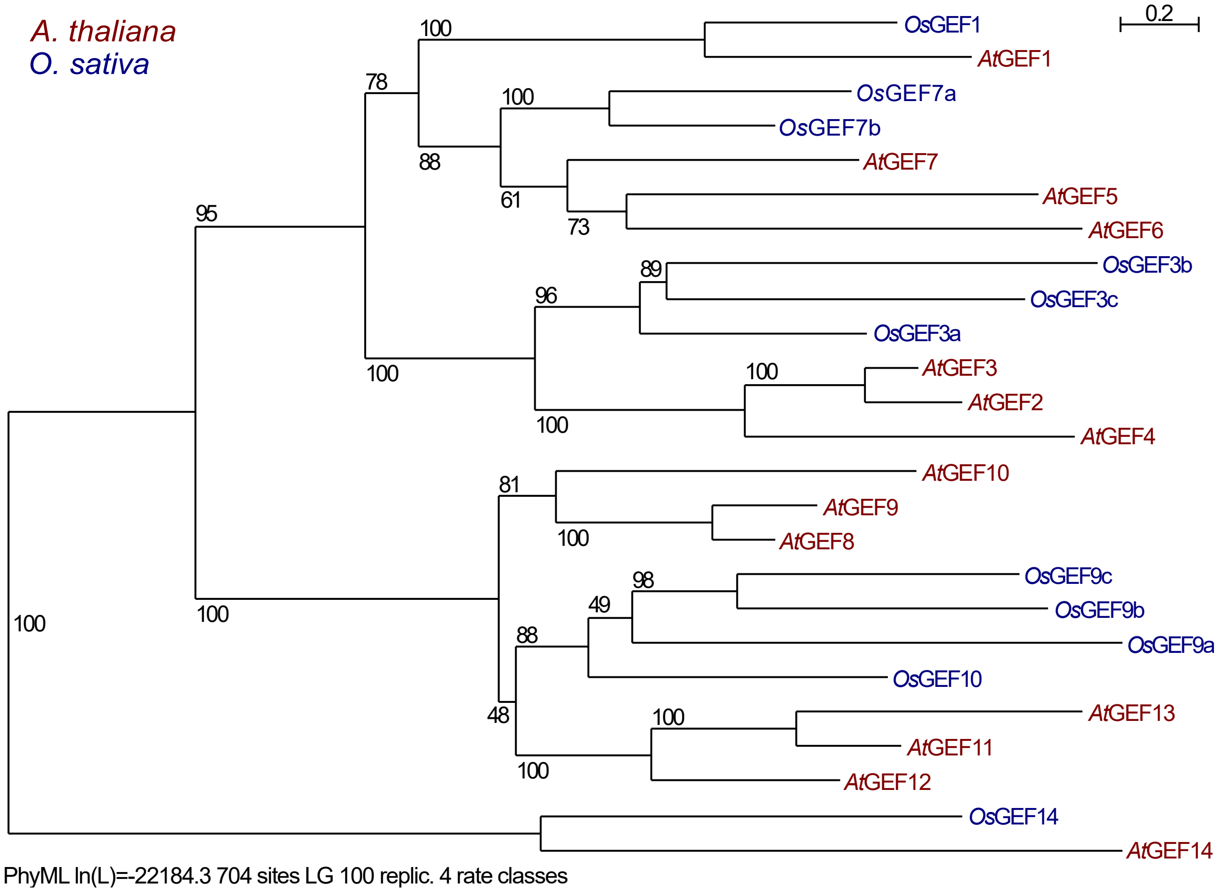


Fig. S1: ***A. thaliana* and *O. sativa* PRONE-GEF**s. Maximum likelihood analysis and annotation of *Os*GEFs based on *A. thaliana* nomenclature in Berken *et al.* (2005).

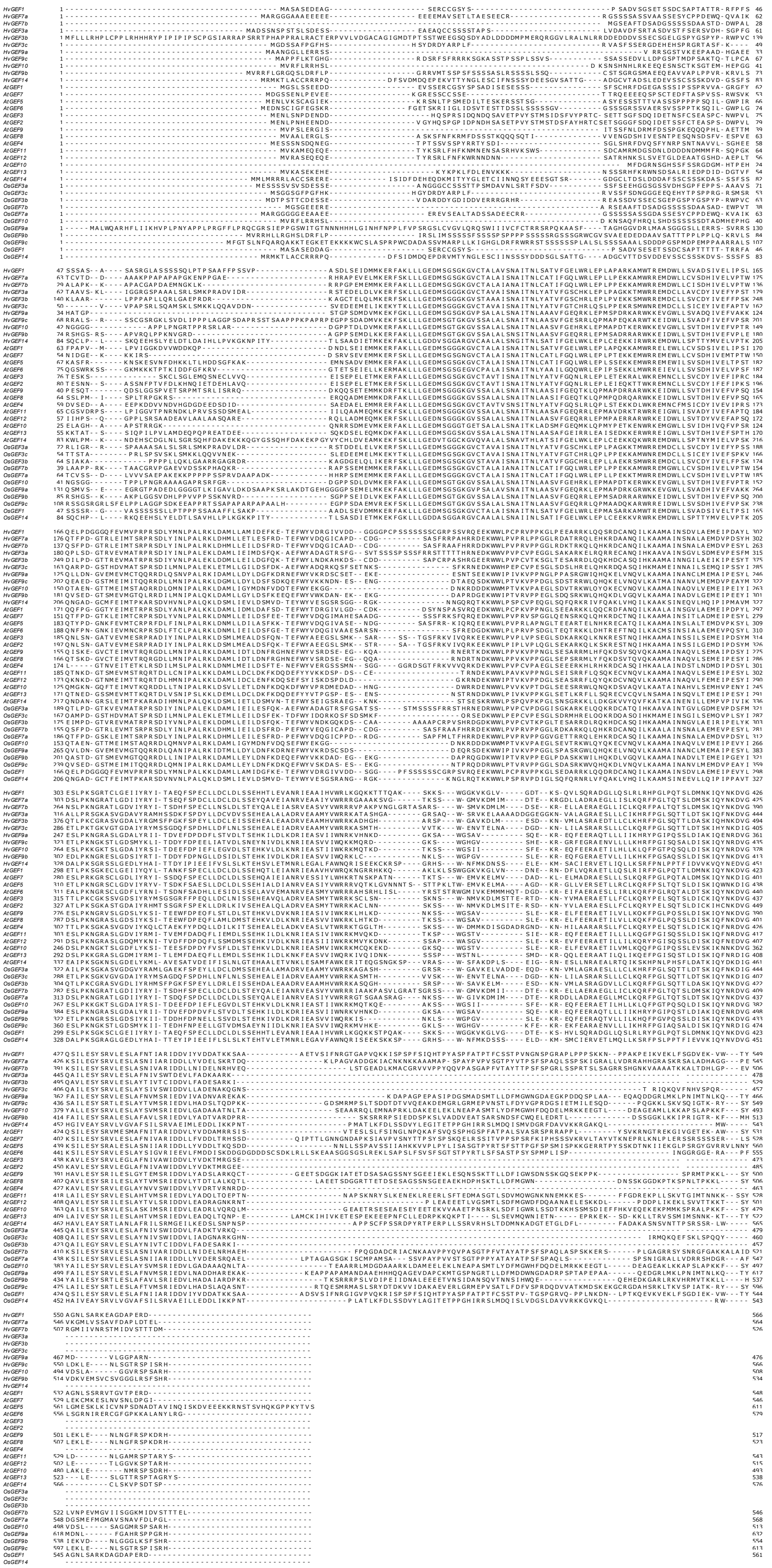

Fig. S2: **MUSCLE alignment of *A. thaliana*, *O. sativa* and *H. vulgare* PRONE-GEFs** as basis for phylogenetic tree in Fig. 1.

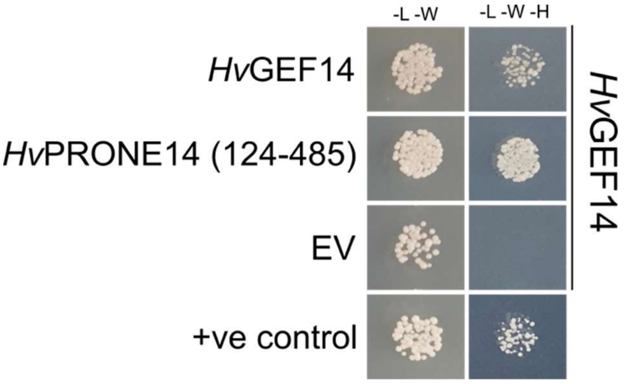


Fig. S3: ***Hv*****GEF14 homo-dimerises with full length and PRONE domain (amino acids 124-485) in yeast-two hybrid.** Protein-protein interaction shown on media containing amino acid mix without leucine (-L), tryptophan (-W), histidine (-H). Growth on –L-W medium shows successful plasmid transformation. Representative image of three experiments with the same result. EV, empty vector; +ve control, positive control.

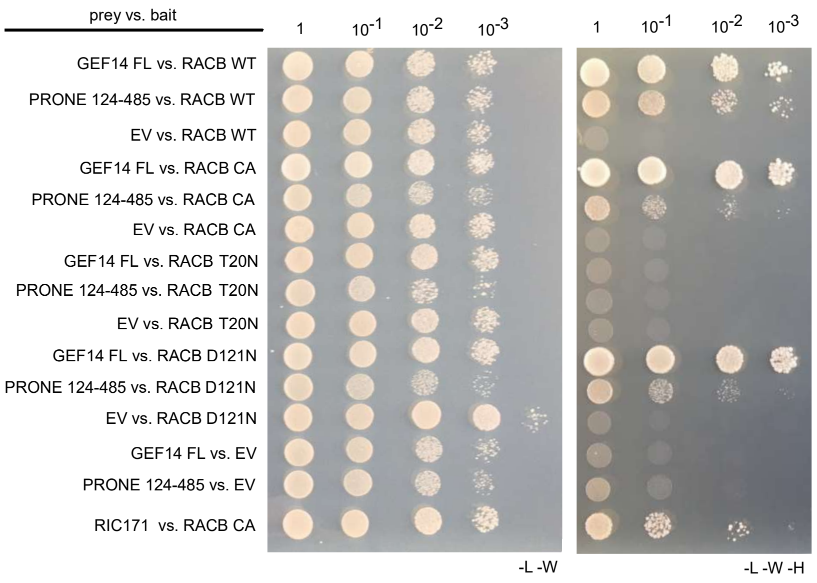


Fig. S4: **Original images of *Hv*GEF14 full length and PRONE domain (amino acids 124-485) interaction with *Hv*RACB variants (wild type, WT; G15V; D121N; T20N) via yeast-two hybrid.** Successful yeast transformation shown on medium containing amino acid mix without leucine (-L) and tryptophan (-W). Medium containing amino acids mix without L, W and histidine (-H) was used as selection for protein-protein interaction. Yeast colonies were grown in liquid -L-W medium and dropped onto plates in a dilution series. Representative image of three experiments with the same result.

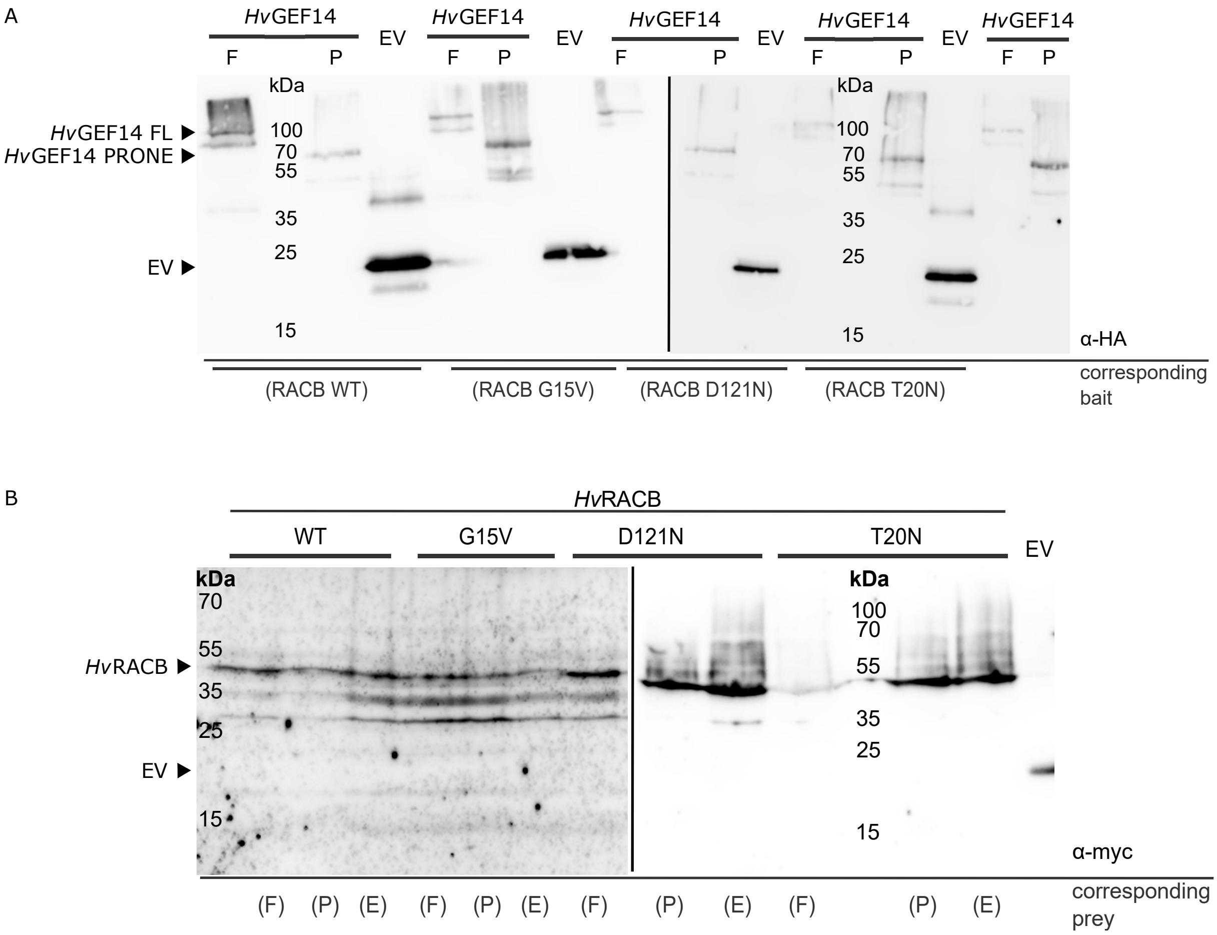


Fig. S5: **Western Blot of yeast proteins on PVDF membranes.** (A) Prey proteins fused to HA-tag and labelled with anti-HA-HRP (3F10, Roche). Detection with DURA chemiluminescence substrate (Thermo Scientific). (B) Bait proteins fused to myc-tag and labelled with c-myc (9E10) and m-IgGκ BP-HRP (sc-516102) antibodies (Santa Cruz Biotechnology). Detection with FEMTO chemiluminescence substrate (Thermo Scientific).

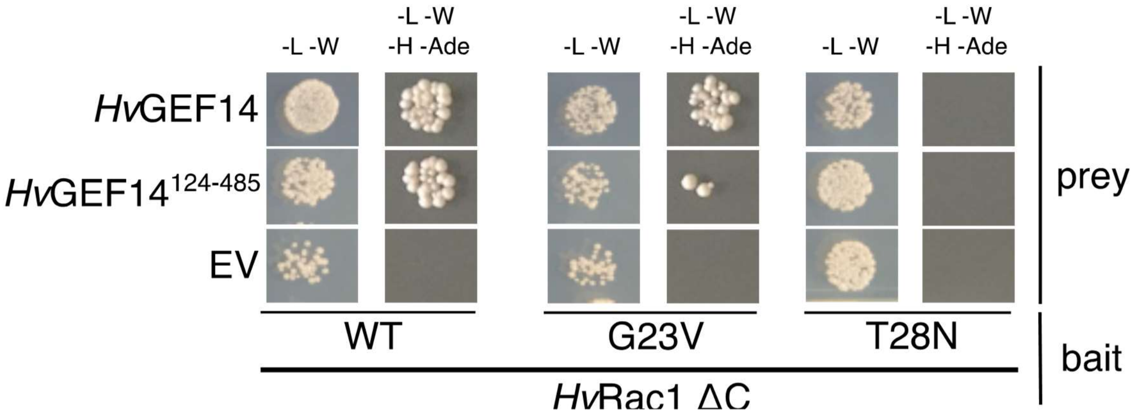


Fig. S6: ***Hv*RAC1 wild type (WT), G23V constitutively activated (CA), T28N dominant negative (DN) variants were tested as bait against prey constructs *Hv*GEF14 full length, *Hv*GEF14 124-485 (PRONE domain) or empty vector (EV).** Interaction of proteins shown on media containing amino acid mix without leucine (L), tryptophan (W), histidine (H), and adenine (Ade) (-L-W-H-Ade) in a dilution series. Representative image of three experiments with the same result.

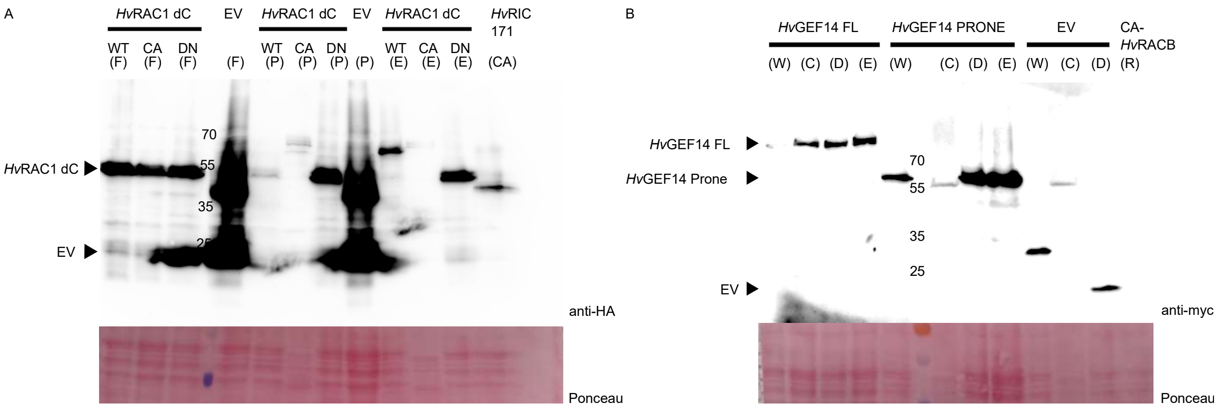


Fig. S7: **Western Blot of yeast proteins on PVDF membranes.** (A) *Hv*GEF14 variants (prey proteins) fused to HA-tag and labelled with anti-HA-HRP (3F10, Roche). Detection with DURA chemiluminescence substrate (Thermo Scientific). (B) *Hv*RAC1 (bait proteins) fused to myc-tag and labelled with c-myc (9E10) and m-IgGκ BP-HRP (sc-516102) antibodies (Santa Cruz Biotechnology). Detection with FEMTO chemiluminescence substrate (Thermo Scientific).

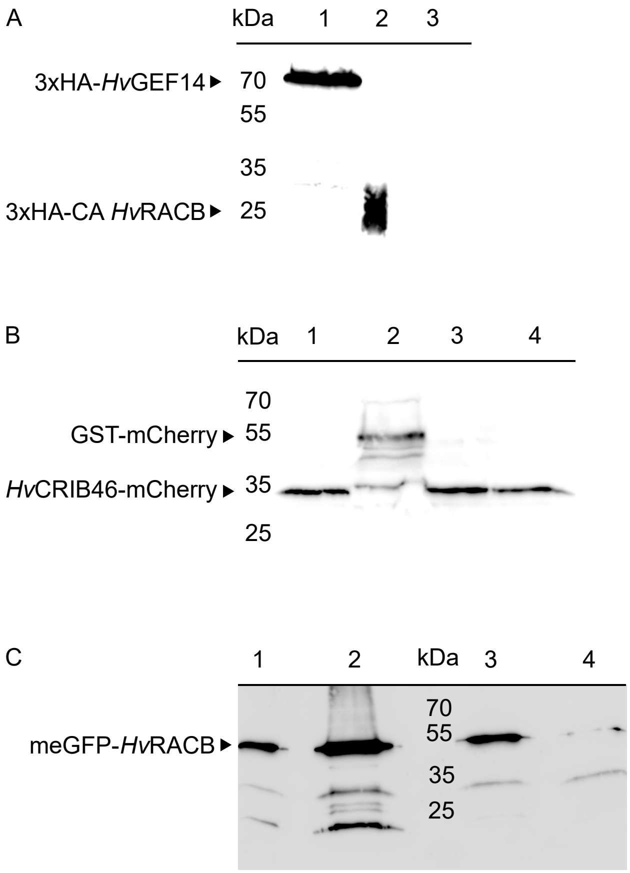


Fig. S8: **Western Blot of proteins extracted from *N. benthamiana* on PVDF membranes.** (A) 3xHA-*Hv*GEF14 co-expressed with meGFP-*Hv*RACB, *Hv*CRIB46-mCherry and P19 in lane 1. 3xHA-CA *Hv*RACB extracted from transgenic barley plants used as positive control in lane 2. Negative control in lane 3 with untagged *Hv*GEF14 co-expressed with meGFP-*Hv*RACB, *Hv*CRIB46-mCherry and P19. Proteins were labelled with anti-HA-HRP (3F10, Roche). (B) *Hv*CRIB46-mCherry co-expressed with meGFP-*Hv*RACB, 3xHA-*Hv*GEF14 and P19 in lane 1. GST-mCherry co-expressed with meGFP-*Hv*RACB and P19 in lane 2. *Hv*CRIB46-mCherry co-expressed with meGFP-*Hv*RACB and P19 in lane 3. *Hv*CRIB46-mCherry co-expressed with meGFP-CA *Hv*RACB and P19 in lane 4. Proteins were labelled with anti-RFP-rat mAb (5F8) and anti-rat (A9542) antibodies (ChromoTek). (C) meGFP-*Hv*RACB co-expressed with *Hv*CRIB46-mCherry, 3xHA-*Hv*GEF14 and P19 in lane 1. meGFP-*Hv*RACB co-expressed with GST-mCherry and P19 in lane 2. meGFP-*Hv*RACB co-expressed with *Hv*CRIB46-mCherry and P19 in lane 3. meGFP-CA *Hv*RACB co-expressed with *Hv*CRIB46-mCherry and P19 in lane 4. Proteins were labelled with anti-GFP (B-2) (sc-9996) and m-IgGκ BP-HRP (sc-516102) antibodies (Santa Cruz Biotechnology). Detection with DURA chemiluminescence substrate (Thermo Scientific).
